# Augmented glucose dependency of autoreactive B cells provides a treatment target for lupus

**DOI:** 10.1101/2022.02.01.475510

**Authors:** John J. Wilson, Jian Wei, Andrea R. Daamen, John D. Sears, Elaine Bechtel, Colleen L. Mayberry, Grace A. Stafford, Lesley Bechtold, Amrie C. Grammer, Peter E. Lipsky, Derry C. Roopenian, Chih-Hao Chang

**Affiliations:** The Jackson Laboratory, Bar Harbor, Maine, ME 04609, USA; School of Basic Medical Sciences, Cheeloo College of Medicine, Shandong University, Shandong 250012, China; AMPEL BioSolutions and the RILITE Research Institute, Charlottesville, VA 22902, USA; Department of Microbiology and Immunology, University of North Carolina, Chapel Hill, NC 27599, USA; Graduate School of Biomedical Sciences and Engineering, University of Maine, Orono, ME 04469, USA; Graduate School of Biomedical Sciences, Tufts University School of Medicine, Boston, MA 02111, USA

## Abstract

Heightened glycolysis is inherent to immune/inflammatory disorders, but little is known of its role in the pathogenesis of systemic lupus erythematosus (lupus). Here, we profile key autoimmune populations in acute and chronic lupus-prone models and their response to glycolytic inhibition. We demonstrate that glycolysis is specifically required for autoreactive germinal center B cells (GCB), but not for T follicular helper cells (Tfh) to survive. This augmented reliance on glucose oxidation to maintain ATP production in pathogenic GCB renders them highly susceptible to oxidative stress-induced apoptosis triggered by glycolysis blockade via 2-deoxyglucose (2DG). We show that 2DG can preferentially reduce GCB in lupus-prone mice, while sparing other autoreactive populations, including Tfh, but still significantly improving lifespan and kidney function. Furthermore, the subset of GCB expressing B-cell maturation antigen (BCMA) exhibits an exaggerated dependence on glycolysis to sustain their growth. Depletion of these cells with a proliferation-inducing ligand-based CAR T-cells leads to greatly prolonged lifespan of mice with severe autoimmune activation. These results reveal that glycolysis dependent GCB, especially those expressing BCMA, are key lupus mediators and highlight that they can be selectively targeted to improve disease outcomes for lupus patients.

## Introduction

The autoimmune disease systemic lupus erythematosus (SLE, lupus) is characterized by loss of self-tolerance, leading to a dysregulated expansion of hyperactivated T and B cells ^1,2^. Underlying lupus is the induction of T-cell-dependent activation and clonal expansion of autoreactive B cells in both germinal centers (GCs) and extra-follicular foci, resulting in their differentiation into plasma cells that secrete pathogenic autoantibodies, which are the primary cause of lupus nephritis ^3,4^. B-cell maturation antigen (BCMA, or TNFRSF17) is particularly upregulated in mature B cells as they differentiate toward plasma cells. Moreover, increasing expression of BCMA has also been observed on lupus B cells, and has been linked with increased activation via interactions with the cytokines TNFSF13 and 13B ^5,6^. B cells and particularly activated GC-like B cells, therefore present an attractive target for treatment of lupus, but to date only belimumab, a monoclonal antibody to TNFSF13B has been approved as a targeted treatment for this autoimmune disorder ^7^.

Autoreactive CD4^+^ T follicular helper cells (Tfh), are also critical drivers of lupus autoimmunity ^8,9^ by engaging follicular B cells in cognate and costimulatory interactions and secreting cytokines, especially interleukin (IL)-21, that promote germinal center B cells (GCB) proliferation and differentiation ^8,10,11^. In turn, the expression of costimulatory molecules, such as ICOS-L and CD80/CD86 on activated B cells in GC further promotes Tfh activation, differentiation, and Bcl6 expression ^12^. Because of the co-dependency of Tfh and GCB, both are critical regulators of spontaneous GC formation during lupus ^2,9^. A primary metabolic adaptation of activated lymphocytes is increased glucose metabolism ^13^, likely mirrored in hyperactive autoimmune Tfh and GCB. This increased glucose metabolism not only provides a rapid source of ATP, but also glycolytic intermediates required for enhanced proliferation and effector functions ^14,15^. With this accentuated glycolytic demand comes a potential avenue for therapeutic intervention by exploiting differential metabolic activity between these cell types for precise treatment of disease. However, the distinct metabolic requirements of discrete autoreactive T and B cells remain unclear.

Here, we demonstrate that activated autoimmune B cells closely resembling GCB (GL7^+^ B cells), but not autoreactive Tfh (CXCR5^+^ PD1^+^ CD4^+^ T cells), exhibit markedly enhanced glycolytic activity that makes them selectively susceptible to therapeutic intervention in spontaneous mouse models of lupus. Compared with Tfh, GCB show a higher glycolysis rate and glucose dependency. We found that 1 week of glycolytic inhibition (short-term) with 2-deoxyglucose (2DG) preferentially reduced GCB through oxidative stress-induced apoptosis, while leaving Tfh numerically and functionally unaffected. This numerical reduction of GCB, significantly reduced mortality rates and protected mice from lupus nephritis. We also found that a subset of GCB expressing BCMA exhibit markedly elevated glycolytic rates and a correspondingly increased sensitivity to glucose inhibition. Reduction of these cells with TNFSF13 (APRIL)-based CAR-T cells significantly delayed lupus progression, suggesting the importance of these cells in the pathogenesis of lupus. Overall, this work serves as a pre-clinical proof-of-principle that metabolic modulation via 2DG preferentially targets and reduces autoreactive GCB, especially the BCMA-expressing subset. Moreover, this reduction is directly linked with reduced lupus severity, revealing an exploitable vulnerability to treat lupus, while minimizing the broad-spectrum immunosuppression.

## Results

### Long-term glycolytic inhibition impacts severe autoimmunity

Long-term glycolytic inhibition via 2DG can attenuate development of cellular disease phenotypes in chronic lupus-prone mice ^16,17^. We profiled autoreactive immune cell populations over the course of disease progression in the acute lupus-prone mouse model, BXSB.Cg-*Cd8a^tm1Mak^ Il15^tm1Imx^ Yaa* (*Yaa* DKO) ^18^, and asked whether long-term glycolytic inhibition could result in increased lifespan. Six-week-old pre-symptomatic *Yaa* DKO mice showed decreased frequencies of circulating CD4^+^ T cells when compared with healthy controls, increased frequencies of circulating Tfh (cTfh), effector T (Teff), CD25^+^ CD4^+^ T cells, and myeloid cells (**Figure 1A** and **Supplemental Figure 1A**). After 4-weeks (long-term) of 2DG treatment, many of the cellular aberrations observed in untreated *Yaa* DKO mice (decreased CD4^+^ T-cell frequencies, increased Teff and cTfh frequencies) were suppressed, whereas frequencies of CD25^+^ CD4^+^ T cells, total B cells, and myeloid cells were unaffected over the 7 weeks of treatment. Moreover, frequencies of B cells that were positive for GL7, a marker of activated B cells in the periphery ^19,20^ and of GCB in the splenic and lymphatic compartments ^17,21^, were reduced to below those of control mice (**Figure 1A** and **Supplemental Figure 1A**). Flow cytometric analysis of spleen cells from untreated symptomatic *Yaa* DKO mice indicated expansion of Teff, Tfh, GCB, myeloid, plasmablasts/plasma, and CD25^+^ CD4^+^ T cells; 7 weeks of 2DG treatment reversed these abnormalities (**Figure 1B** and **Supplemental Figure 1B**). 2DG-treated mice showed complete survival with the first fatality in the treated group occurring ∼10 weeks after 2DG withdrawal, by which time 95% of untreated mice had died (**Figure 1C**). Notably, the significant reduction of peripheral GL7^+^ B cells was mirrored in splenic GCB, suggesting that long-term glycolytic inhibition had a comparable effect on GL7^+^ B cells in both compartments. Overall, these results indicate that glycolytic inhibition via 2DG is effective in ameliorating autoimmune activation and increasing survival in the model of severe lupus.

**Figure 1.**
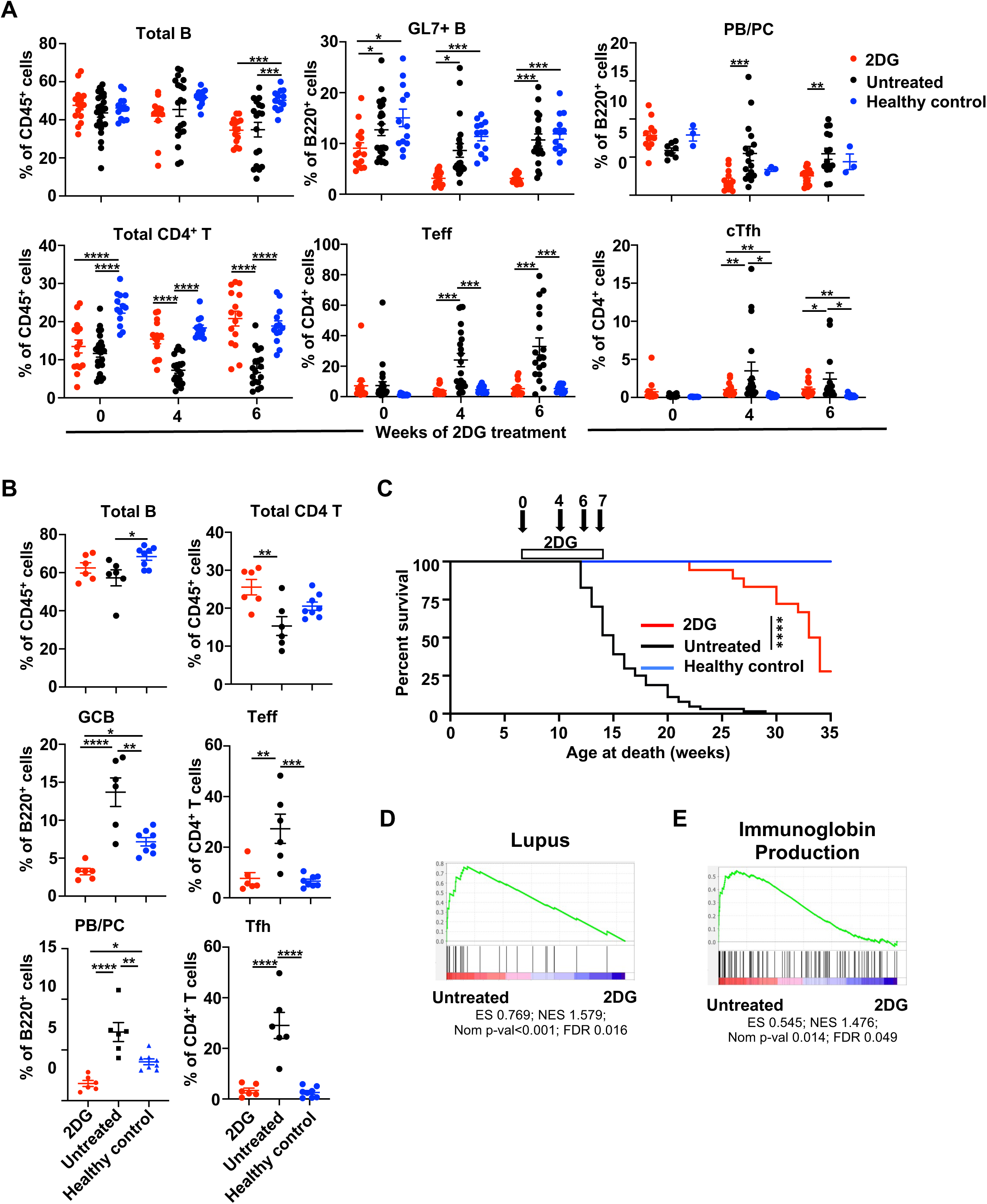
Glycolytic inhibition decreases disease biomarkers and increases life expectancy in lupus-prone mice. (**A**) Flow results of circulating lymphocytes of *Yaa* DKO mice treated with 2DG at 6 weeks of age for indicated weeks or untreated, and healthy controls (n=15-25 mice per group). (**B**) Flow results of splenic T and B cells of 13-week old *Yaa* DKO mice treated with 2DG for 7 weeks or untreated, or untreated healthy controls (n=6-8 mice per group). PB/PC, plasmablasts/plasma cells. (**C**) Survival of *Yaa* DKO mice treated with 2DG at 7 weeks of age, or untreated and healthy controls, with arrows showing sampling times (**A**) (n=15-25 mice per group). Data are from at least three independent experiments. (**D**-**E**) GSEA plots of splenocytes from 10-week-old *Yaa* DKO mice untreated vs. treated with 2DG for 4 weeks (n=4 mice per group). Enrichment of gene sets in (**D**) lupus and (**E**) immunoglobulin production. Each dot represents one mouse. Error bars represent mean±SEM; \**p*<0.05, \*\**p*<0.01, \*\*\**p*< 0.001, \*\*\*\**p*<0.0001 using one-way ANOVA with Bonferroni’s multiple comparison tests (**A**-**B**) or with Mantel-Cox test (**C**).

### Long-term glycolytic inhibition potently affects B-lineage cells

Since the gene-expression patterns of T and B cells from lupus patients differ with those from healthy individuals ^22,23^, we investigated whether 2DG treatment alters the dysregulated gene expression patterns in *Yaa* DKO mice. Initially, we carried out Gene Set Enrichment Analysis (GSEA) ^24,25^ to compare gene-expression profiles of spleen cells from long-term 2DG-treated and untreated *Yaa* DKO mice. Many genes dysregulated in human lupus were enriched in untreated mice, including those involved in immunoglobulin production (**Figure 1, D and E**). To examine the effect of 2DG treatment in greater detail, we carried out Gene Set Variation Analysis (GSVA) ^26^ with a panel of curated gene modules and compared transcriptomic signatures in spleens between 2DG-treated and -untreated symptomatic *Yaa* DKO mice at 10 weeks of age, and untreated pre-symptomatic mice at 6 weeks of age (**Figure 2A**). GSVA scores indicative of activated B cells and plasma cells were significantly decreased with 2DG treatment. However, gene signatures of total T cells, total B cells, activated T cells and myeloid cells were unaffected (**Figure 2A**). An analysis of metabolic pathways revealed that 2DG treatment decreased expression of genes controlling glycolysis and glutamine metabolism (**Figure 2B**).

**Figure 2.**
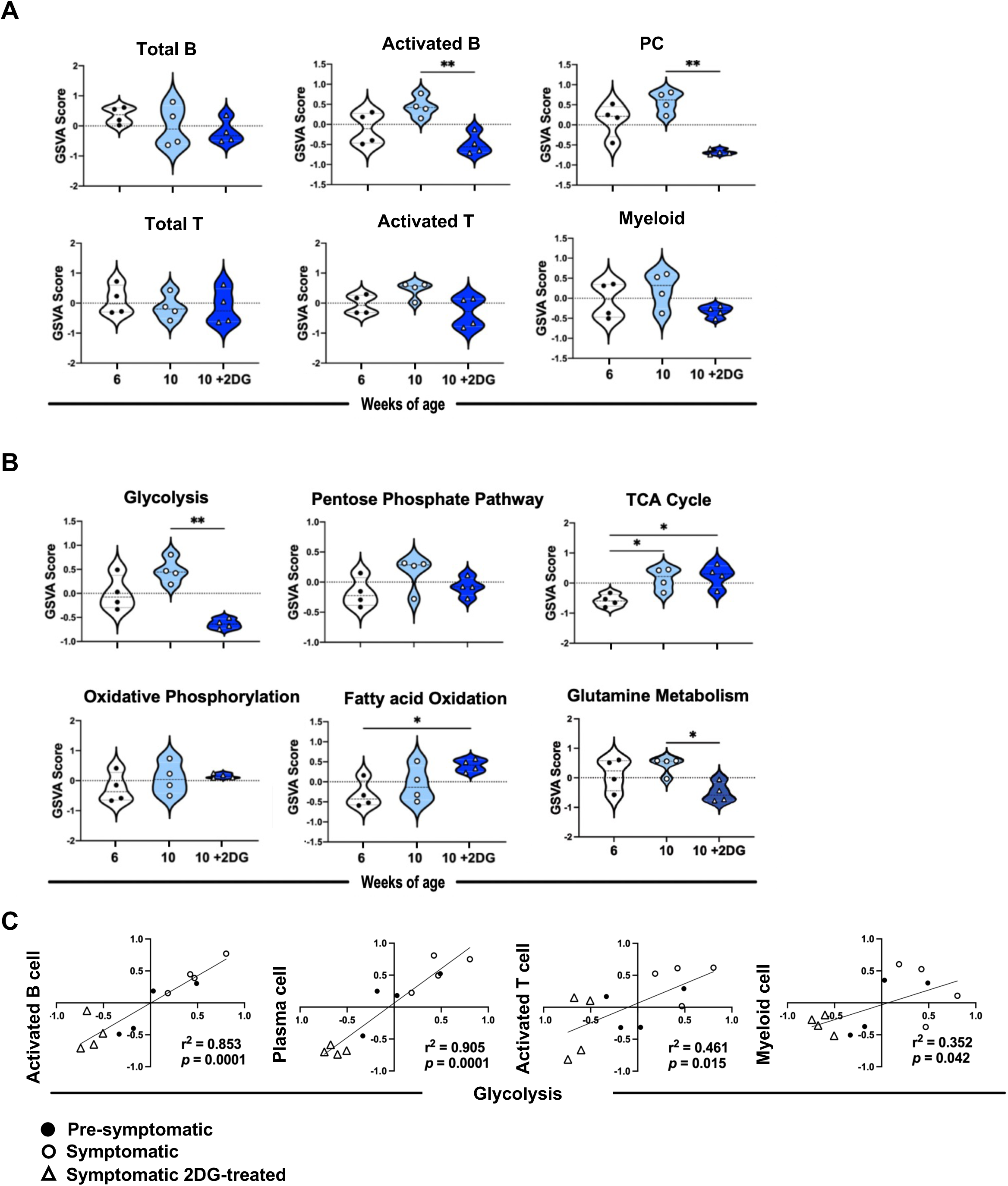
Long-term glycolytic inhibition has a selectively potent effect on gene signatures of B cells. Violin plots of gene-expression data from splenocytes of individual 6-week-old (pre-symptomatic) untreated *Yaa* DKO mice (●), or 10-week-old *Yaa* DKO mice treated with 2DG (Symptomatic 2DG-treated) for 4 weeks (△) or untreated (Symptomatic) (○), were analyzed by GSVA for enrichment of (**A**) immune cell and (**B**) metabolic pathway gene signatures. (n=4 mice per group). \**p*<0.05, \*\**p*<0.01 using Brown Forsythe and Welch’s ANOVA with Dunnett’s T3 multiple comparison tests. **(C)** Linear regression between GSVA scores for glycolysis gene signatures (n=4 mice per group). The goodness of fit for each comparison is displayed as the r^2^ value and the slope of the regression line is displayed as the p-value. Correlations with p<0.05 were considered significant. PC, plasma cells.

Linear regression analyses indicated a strong positive correlation between the glycolysis and cellular signatures for activated B and plasma cells, and far weaker positive correlations for activated T cells and myeloid cells (**Figure 2C**). No cellular signatures were correlated with TCA cycle or oxidative phosphorylation, whereas plasma cells had a strong negative correlation with oxidative phosphorylation (OXPHOS). Both activated B cells and plasma cells had strong positive correlations with glutamine metabolism, whereas myeloid cells had a much weaker association (**Supplemental Figure 2**). Notably, although the activated T cell gene signature was also influenced, it was decreased to a much lesser degree. These data indicate that 2DG-mediated glycolytic inhibition preferentially influences metabolic regulation of activated B cell populations.

### Autoreactive GCB show elevated glucose dependency compared to Tfh

Gene expression analysis suggested that activated B lineage cells appear to be more sensitive to long-term glycolytic inhibition than T cells, consistent with the greater depletion of GCB (to below normal levels) than Tfh (**Figure 1B**). Given this disparity, we hypothesized that the metabolic requirements of autoreactive GCB and Tfh might differ and, specifically, that GCB might be more dependent on glycolysis. To test this, splenic GCB and Tfh from symptomatic *Yaa* DKO mice were sorted to assess their bioenergetic profiles. Autoreactive GCB were found to have higher extra-cellular acidification rates (ECAR), an indicator of glycolysis, and oxygen consumption rates (OCR), an indicator of mitochondrial OXPHOS, compared to Tfh (**Figure 3A**), and other effector T and non-GCB cells (**Figure 3B**). This high bioenergetic profile was also found in GCB isolated from symptomatic NZBWF1 late-onset lupus-prone mice (**Supplemental Figure 3A**). To determine whether this elevated metabolic demand was disease-related, basal ECAR and OCR in GCB and Tfh from healthy, pre-symptomatic and symptomatic *Yaa* DKO mice were assessed. We found that symptomatic mice displayed significantly higher ECAR in both GCB and Tfh, but that only GCB showed elevated OCR compared to pre-symptomatic or healthy mice (**Figure 3C**). Autoreactive GCB also showed greater expression of the major glucose transporter Glut1 (**Figure 3D**), and significantly higher 2-NBDG uptake than did Tfh (**Figure 3E**). In addition, GCB, cultured *ex vivo*, reduced the concentration of glucose in the medium to a significantly greater degree than Tfh (**Figure 3F**). Together, these data demonstrate that GCB exhibit a higher glucose usage and glycolysis rate than do Tfh.

**Figure 3.**
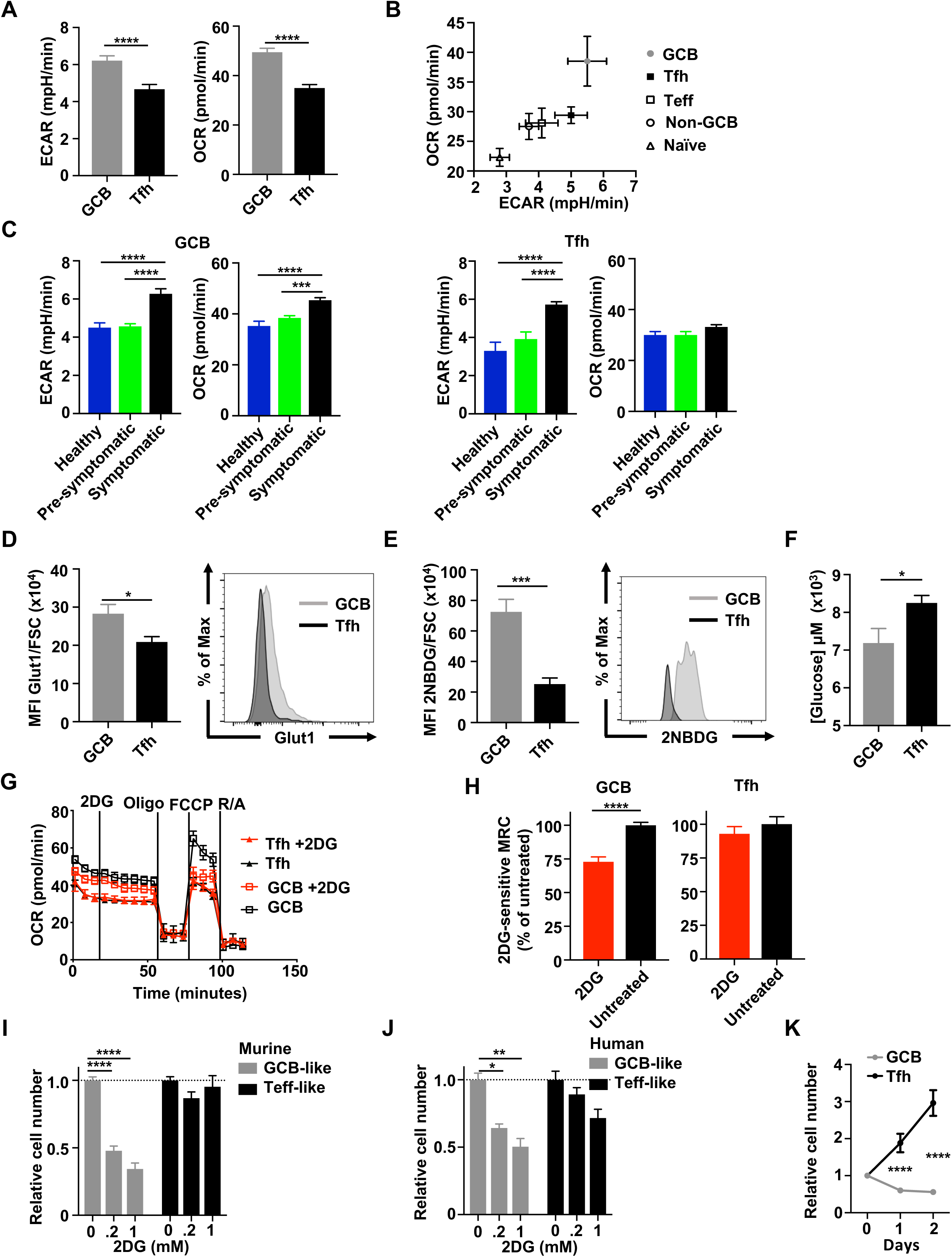
Autoreactive GBC display augmented glycolytic dependency over Tfh. (**A-I, K**) Bioenergetic analysis of spleen cells from symptomatic *Yaa* DKO mice. (**A**) Basal ECAR and OCR of GCB and Tfh. (**B**) Basal OCR versus ECAR of *Yaa* DKO GCB (●), Tfh (▪), Teff (□), non-GCB (○) and naïve C57BL/6 splenocytes (Δ). (**C**) Basal ECAR and OCR of GCB and Tfh from healthy, pre-symptomatic and symptomatic mice. (**D**) Glut1 expression on GCB and Tfh, normalized to cell size (FSC). (n=4-5 mice per group). (**E**) *In vivo* glucose uptake in GCB and Tfh measured as 2-NBDG mean fluorescence intensity (MFI) normalized to FSC (n=4-5 mice per group). (**F**) Glucose concentration remaining after culture of Tfh or GCB in complete medium for 24 hours, normalized to live-cell numbers. (**G**) OCR and MRC measurements of GCB and Tfh, with or without 2DG before treatment with oligomycin (Oligo), FCCP, and rotenone/antimycin (R/A). (**H**) 2DG-effect on MRC was normalized to untreated cells and assessed as the contribution of glucose metabolism to OXPHOS (2DG-sensitive MRC) in GCB and Tfh following FCCP exposure. Relative live-cell numbers of (**I**) murine or (**J**) human GCB-like or Teff-like cells after 24-hour treatment with 2DG, normalized to untreated controls. (**K**) Live GCB and Tfh numbers in the presence of 1 mM 2DG, normalized to untreated controls. Data are from at least three independent samples. Error bars represent mean±SEM; \**p*<0.05, \*\**p*<0.01, \*\*\**p*<0.001, \*\*\*\**p*<0.0001 using one-way ANOVA with unpaired two-tailed Student’s *t*-tests (**A**,**D**-**F,H**) or Bonferroni’s multiple comparison tests (**C**,**I**-**K**).

We next investigated whether enhanced glycolysis supports OXPHOS, by treating autoreactive GCB and Tfh with 2DG in the presence of glucose. Although 2DG reduced the basal ECAR of both GCB and Tfh to similar levels (**Supplemental Figure 3B**), 2DG-sensitive maximal respiratory capacity (MRC) was significantly decreased (27%) in GCB, but not Tfh, non-GCB, nor in Teff (**Figure 3, G** and **H**, and **Supplemental Figure 3C**), after FCCP-mediated mitochondrial decoupling, an indicator of OXPHOS reliance on a nutrient substrate fueling mitochondria ^27^. A similar 2DG sensitivity was also found in GCB (30%) from NZBWF1 mice (**Supplemental Figure 3D**). These results suggest that 2DG-mediated glycolytic inhibition selectively impacts OXPHOS in GCB. To confirm whether fueling the mitochondrial matrix in GCB depends on glycolysis, we exposed cells to UK5099, which blocks the import of glycolysis-derived pyruvate into mitochondria. Treatment with UK5099 significantly impaired the MRC of GCB (15%) but had no effect on Tfh (**Supplemental Figure 3E**). We further evaluated whether mitochondrial fueling of GCB also depends on other catabolic pathways.

Treatment of GCB and Tfh with BPTES, a glutaminase inhibitor ^28^; etomoxir, which suppresses mitochondrial fatty acid oxidation (FAO) or thioridazine, a selective inhibitor of peroxisomal FAO ^29^; resulted in little to no reduction in basal OCR in either cell type (**Supplemental Figure 3, F**–**H**). Treatment with BPTES resulted in a small but significant reduction of MRC (7.7%) in GCB but not in Tfh (**Supplemental Figure 3I**); however, this reduction was much less than that found with glycolytic inhibition (**Figure 3H**). Conversely, no change in MRC was observed in either GCB or Tfh treated with etomoxir or thioridazine (**Supplemental Figure 3, J** and **K**). Together, these results are consistent with the mechanism of GCB reliance on glucose as the primary anaplerotic precursor to support mitochondrial function, whereas autoreactive Tfh exhibit greater metabolic flexibility.

To test whether glycolytic inhibition preferentially affects GCB survival, mouse splenocytes were activated *in vitro*. Numbers of differentiated GCB-like cells were significantly decreased by 2DG exposure, whereas Teff-like cells were unaffected (**Figure 3I**). A similar trend in 2DG sensitivity was found in activated human PBMCs (**Figure 3J**). Glycolytic blockade with the hexokinase inhibitor 3-bromopyruvate also selectively decreased numbers of murine GCB-like cells (**Supplemental Figure 3L**), further confirming the efficacy of glycolytic inhibition. We next assessed the effect of 2DG treatment on the survival of splenic GCB and Tfh isolated from symptomatic *Yaa* DKO mice. One-day 2DG exposure significantly reduced GCB numbers, which decreased further after two days, whereas expansion of Tfh was observed (**Figure 3K**). These data show that 2DG treatment preferentially affects survival of highly-glycolytic GCB, while sparing the less bioenergetic Tfh.

### Short-term glycolytic inhibition markedly reduces the numbers of GCB but not Tfh in diseased lupus-prone mice

While early long-term 2DG treatment was effective at preventing disease in *Yaa* DKO mice, the effects of prolonged glycolytic inhibition might be too oppressive for clinical application. Given their increased glycolytic dependency, we hypothesized that short-term 2DG treatment might selectively inhibit GCB *in vivo*. First, we assessed circulating activated B cells and Tfh in symptomatic *Yaa* DKO mice as treatment progressed and found that GL7^+^ B cell numbers dropped significantly by day 7 of therapy with no cTfh reduction (**Figure 4A**). No other circulating populations showed significantly reduced numbers compared to untreated controls with this 1-week (short-term) treatment course (**Figure 4B** and **Supplemental Figure 4A**). Next, a similar reduction in GCB was observed in lymph nodes (**Supplemental Figure 4B**) and spleens (**Figure 4B** and **Supplemental Figure 4C**) of the mice, with no significant numerical reductions in other populations. Numbers of splenic follicular regulatory T (Tfr), Treg, and transitional, follicular, and marginal zone B cells, were also unaffected following short-term treatment (**Supplemental Figure 4, D**–**F**). 2DG-induced reduction of cell numbers was likewise observed in B220^+^ CD95^hi^ CD38^low^ gated GCB ^29^ (**Supplemental Figure 4G**). Moreover, symptomatic NZBWF1 mice treated with short-term 2DG displayed reductions in both circulating GL7^+^ B cells and splenic GCB numbers, similar to those in *Yaa* DKO mice (**Figure 4C**). These data suggest that increased sensitivity to glycolytic inhibition is a uniform feature of autoreactive GCB, rendering them comparatively more vulnerable than other autoimmune cell types.

**Figure 4.**
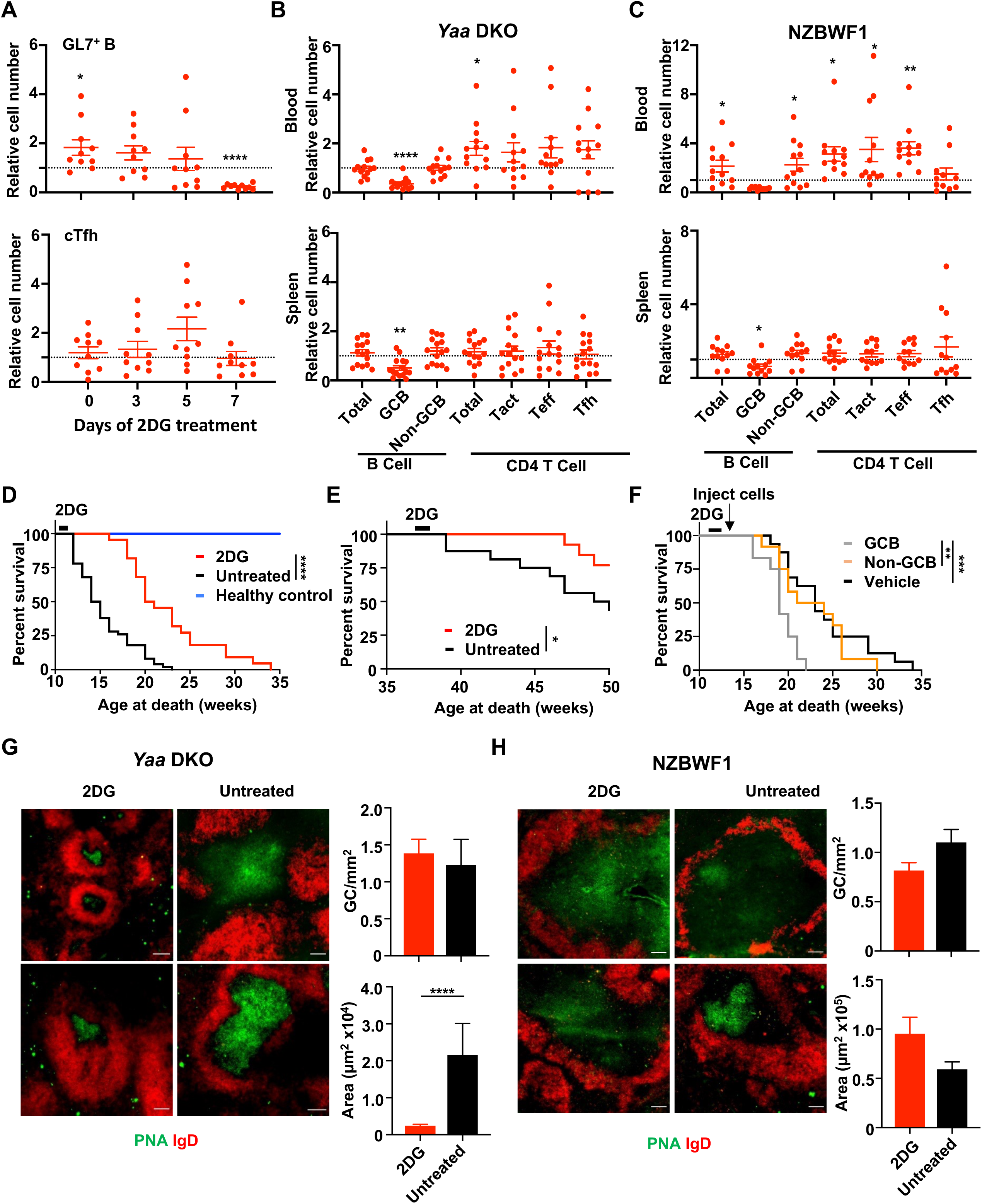
Short-term glycolytic inhibition selectively reduces autoreactive GCB and improves survival outcomes. (**A**) Flow plots of relative numbers of circulating cells from *Yaa* DKO mice treated with 2DG at days 0, 3, 5 and 7, normalized to untreated controls (n=10). (**B**-**C**) Cell numbers from peripheral blood and spleen normalized to untreated mice after 1 week of 2DG exposure in (**B**) *Yaa* DKO (n=13-15 mice per group) and (**C**) in NZBWF1 mice (n=12 mice per group). (**D**) Survival of symptomatic *Yaa* DKO mice treated with 2DG for 1 week starting at 11-weeks-old, age-matched untreated mice, and healthy controls (n=22-50 mice per group). (**E**) Survival of NZBWF1 mice treated with 2DG for 1 week starting at 36-weeks-old and untreated controls (n=13-16 mice per group). (**F**) Survival of 1-week 2DG-treated, symptomatic *Yaa* DKO mice after adoptive transfer of GCB or non-GCB cells (n=14 mice per group). Immunohistochemistry images of (**G**) symptomatic *Yaa* DKO or (**H**) NZBWF1 spleens from mice treated with 2DG for 1 week and untreated control mice were stained with anti-IgD (red) and anti-PNA (green) for germinal center identification. (Scale bars = 100 μm, n=3). Data are from at least two independent experiments. Each dot represents one mouse. Error bars represent mean±SEM; \**p*<0.05, \*\**p*<0.01, \*\*\**p*<0.001, \*\*\*\**p*<0.0001 using one-way ANOVA with Bonferroni’s multiple comparison tests (**A**-**C**), with Mantel-Cox tests (**D**-**F**) or using an unpaired two-tailed Student’s *t*-test (**G-H**).

To test whether this GCB reduction can reverse autoimmune pathology in mice with severe clinical disease, we treated symptomatic *Yaa* DKO mice with 2DG for 1 week and monitored their survival. The mean lifespan of untreated mice was 15.4 weeks, in contrast to 24.1 weeks for 2DG-treated mice (**Figure 4D**). Of note, mice showed no significant decrease in frequencies of autoreactive Tfh and ICOS^+^ T cells after 2DG removal (**Supplemental Figure 4H**), suggesting that therapeutic efficacy was unlikely to be conveyed by a post-treatment reduction of these populations. NZBWF1 mice experience progressive proteinuria and death from lupus nephritis caused by cumulative kidney damage owing to immune complex deposition ^30,31^. Lifespan extension (**Figure 4E**), paired with persistently reduced proteinuria (**Supplemental Figure 4I**), was also observed in NZBWF1 mice treated with 1-week 2DG. These data suggest that diverse murine lupus models can be positively impacted by this short-term 2DG treatment. To confirm the involvement of 2DG-responsive GCB in disease pathogenesis, an adoptive transfer approach was undertaken. As noted above, GCB numbers were reduced in *Yaa* DKO mice with 1-week of 2DG (**Figure 4B**). Following treatment, the mice underwent adoptive transfer of flow-sorted autologous GCB or non-GCB cells from symptomatic *Yaa* DKO mice. The mean lifespans of GCB-recipient mice were significantly shorter (19.0 weeks) than those receiving non-GCB cells (23.8 weeks) or no cells (24.1 weeks) (**Figure 4F**). These data demonstrate that, short-term 2DG treatment was remarkably effective in reversing disease trajectory with a concomitant decrease in lupus nephritis and mortality. We then examined the effect of short-term 2DG treatment on splenic morphology. Histologic examination of spleens showed that treatment of *Yaa* DKO mice with 2DG significantly reduced the size but not the number of GCs (**Figure 4G**). The effects were different in NZBWF1 mice with no significant effect on splenic GC morphology (**Figure 4H**), consistent with the lower efficacy of 2DG in reducing GCB numbers in NZBWF1 mice (**Figure 4, B** and **C**) and implying a need to optimize 2DG treatment in these mice. Taken together, these results demonstrate that autoreactive GCB, shown to be extremely 2DG-sensitive, are highly effective at accelerating lupus.

### Glycolytic inhibition results in preferential induction of apoptosis in GCB

Inhibition of glycolysis can cause cell death through apoptosis ^32^. To determine the mechanism by which autoreactive GCB are reduced, we treated symptomatic *Yaa* DKO mice with 2DG to assess cellular apoptosis. By day 4 of treatment, there was a significant increase in relative frequencies of pre-apoptotic (Annexin^+^/PI^-^) GL7^+^ B cells but not cTfh in 2DG-treated mice (**Figure 5A** and **Supplemental Figure 5A**). With 2DG treatment, there was a trend toward increased apoptotic GL7^+^ B cells on day 7, but no such trend for cTfh (**Figure 5A**). Splenic GCB isolated from 7-day 2DG-treated mice showed increased pre-apoptosis (**Figure 5B**). In Tfh, frequencies of pre-apoptosis were unchanged, whereas apoptosis was significantly decreased (**Figure 5B**). The reduction in GCB numbers could not be explained by a loss of GL7 protein as 2DG did not affect GL7 expression in GCB from *Yaa* DKO mice *in vitro* (**Supplemental Figure 5B**). Moreover, numbers and frequencies of splenic GL7-expressing Teff in *Yaa* DKO mice were unaltered with 2DG treatment (**Supplemental Figure 5C**). Together, these data suggest that short-term 2DG exposure selectively induces apoptosis in autoreactive GCB.

**Figure 5.**
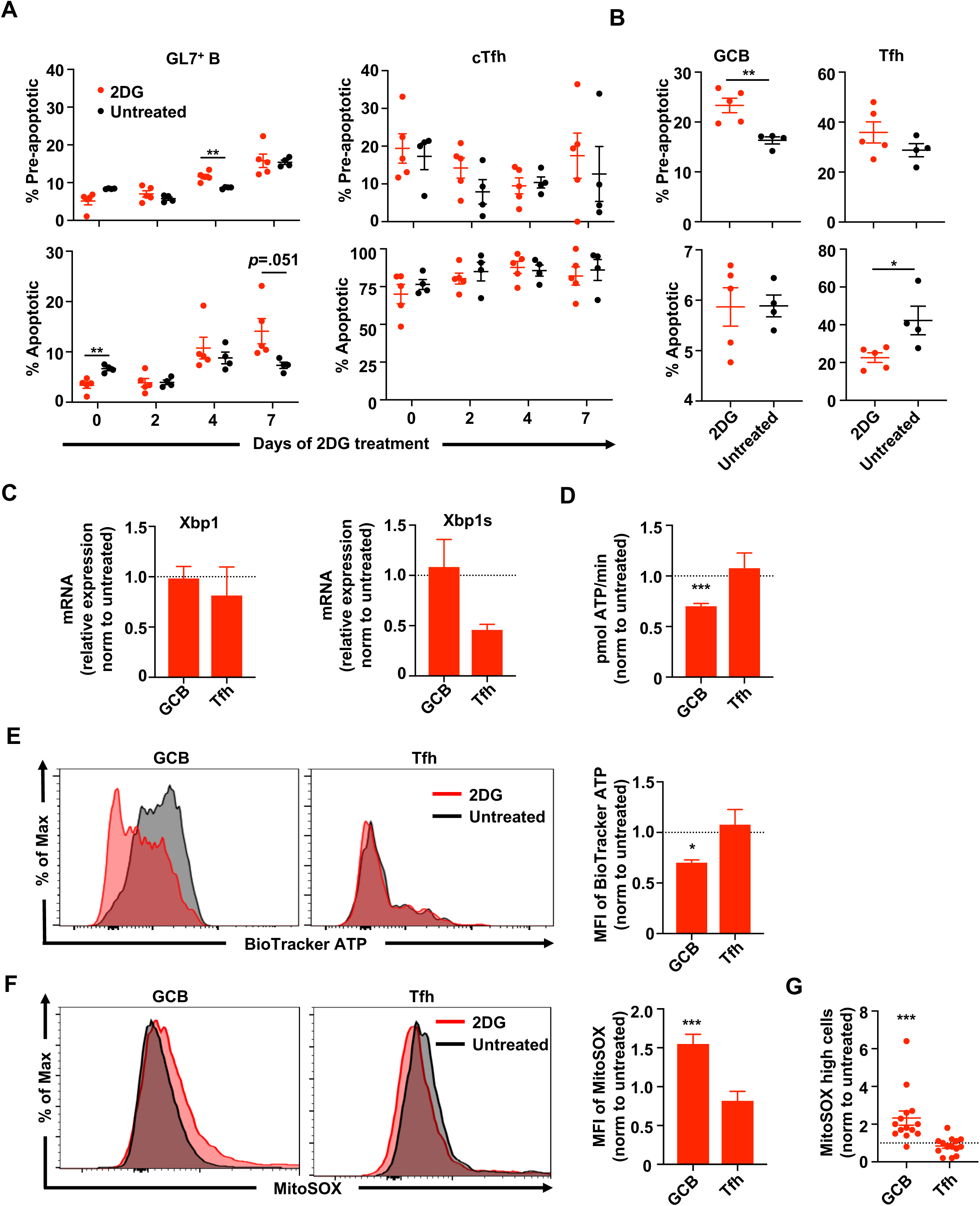
Glycolytic inhibition selectively triggers apoptotic death in GCB. Frequencies of apoptotic and pre-apoptotic cells in (**A**) blood and (**B**) spleens of *Yaa* DKO mice treated with 2DG for 1 week (n=4-5 mice per group). (**C**) Relative *Xbp1* and *Xbp1s* expression on GCB and Tfh from *Yaa* DKO mice treated with 2DG for 1 week, normalized to untreated controls. (n= 4-6 mice per group) (**D**) ATP production rate of GCB and Tfh from *Yaa* DKO mice after 2DG treatment. (**E**) ATP concentration in GCB and Tfh from *Yaa* DKO mice treated with 2DG. (**F**) MFI and (**G**) relative frequency of mitochondrial ROS (MitoSOX) in splenic GCB and Tfh of *Yaa* DKO mice treated with 2DG for 1 week normalized to untreated controls (n=14). Data are from at least three independent samples. Each dot represents one mouse. Error bars represent mean ± SEM; \**p*<0.05, \*\**p*<0.01, \*\*\**p*<0.001 using one-way ANOVA with Bonferroni’s multiple comparison tests (**A**) or unpaired two-tailed Student’s *t*-tests (**B**-**G**).

Apoptosis can be induced by endoplasmic reticulum (ER) stress and the unfolded protein response (UPR) ^32,33^. The inositol-requiring enzyme 1/X-box-binding protein 1 (XBP1) pathway is a potent UPR signaling pathway in mammalian cells, promoting ER-induced apoptosis. To ascertain whether UPR signaling is elevated with 2DG treatment, we assessed mRNA expression of *Xbp1* and its spliced form, *Xbp1s*, as markers of ER stress ^33,34^. *Xbp1* expression in both autoreactive GCB and Tfh were unchanged (**Figure 5C**), whereas expression of *Xbp1s* was decreased in Tfh with 2DG-treatment, suggesting that neither GCB nor Tfh experienced ER stress *in situ* following transient 2DG. Glucose deprivation may reduce cellular ATP production and increase reactive oxygen species (ROS) levels, leading to oxidative stress and the subsequent death of tumor cells ^35,36^. Hence, we interrogated 2DG-treated cells and found that the overall ATP production rate was reduced in autoreactive GCB but not in Tfh, compared with those isolated from untreated mice (**Figure 5D**). This was further confirmed with lower levels of total ATP measured in GCB treated with 2DG compared to untreated controls (**Figure 5E**). We therefore examined the accumulation of mitochondrial ROS as an indicator of oxidative stress ^35^, in *Yaa* DKO mice. Following 2DG treatment, ROS production was significantly elevated in GCB, but was unchanged in Tfh (**Figure 5F**), non-GCB, and Teff (**Supplemental Figure 5D**); whereas neither Tfh nor GCB experienced changes in mitochondrial mass (**Supplemental Figure 5E**). Excessive ROS production in GCB is also linked to increased mitochondrial membrane potential (**Supplemental Figure 5F**). Taken together, these data suggest that the underlying mechanism of selective autoreactive GCB reduction with short-term 2DG exposure was a pro-apoptotic effect from energy deprivation through increased mitochondrial oxidative stress.

### Autoreactive Tfh are not functionally impaired by short-term glycolytic inhibition

Tfh are vital to the formation and stability of the GC, as they provide survival and differentiation signals to GCB ^2,9^. Although numbers of autoreactive Tfh were unaffected by short-term glycolytic inhibition, it remained possible that this treatment impaired Tfh functionality and thus contributed to GCB apoptosis. To test this possible mechanism of reduction, splenic Tfh were isolated from *Yaa* DKO mice following 1-week 2DG treatment and examined for functional markers. Tfh-derived IL-21 is thought to play a pivotal role in lupus pathogenesis and is the major cytokine inducing B-cell differentiation ^10,11,18^. Notably, neither the frequency of IL-21-producing Tfh nor IL-21 expression differ following this treatment (**Figure 6A**). The transcription factor Bcl6 is essential for development of both Tfh and GCB ^37,38^. We found that Bcl6 protein expression was not impacted by 2DG in either Tfh or GCB (**Figure 6B**). In addition, the expression and frequency of Ki-67, a proliferation marker, in both Tfh and GCB were unaffected by 2DG (**Figure 6C**). Furthermore, no treatment-induced reduction in the frequency or expression of the GCB-Tfh interaction molecules ICOS-L, CD40, CD80 and CD86 on GCB (**Supplemental Figure 6A**), or ICOS, CD40L and CD28 on Tfh was observed (**Supplemental Figure 6B**). The basal OCR of sorted Tfh was significantly elevated in mice treated with 2DG compared with untreated mice (**Figure 6D**), suggesting Tfh can better adapt their metabolism to survive glycolytic inhibition. Although the basal ECAR and OCR of GCB were unaffected (**Figure 6D**), the reliance of remaining GCB from 2DG-treated mice on glycolysis became negligible (**Figure 6E**). Together, these results suggest that Tfh and GCB remain functionally intact following short-term glycolytic inhibition, and that this inhibition specifically affects hyperactive GCB with high glycolytic dependency. Overall, these data indicate that the effect of 2DG on the numerical reduction of GCB is direct and not mediated by Tfh.

**Figure 6.**
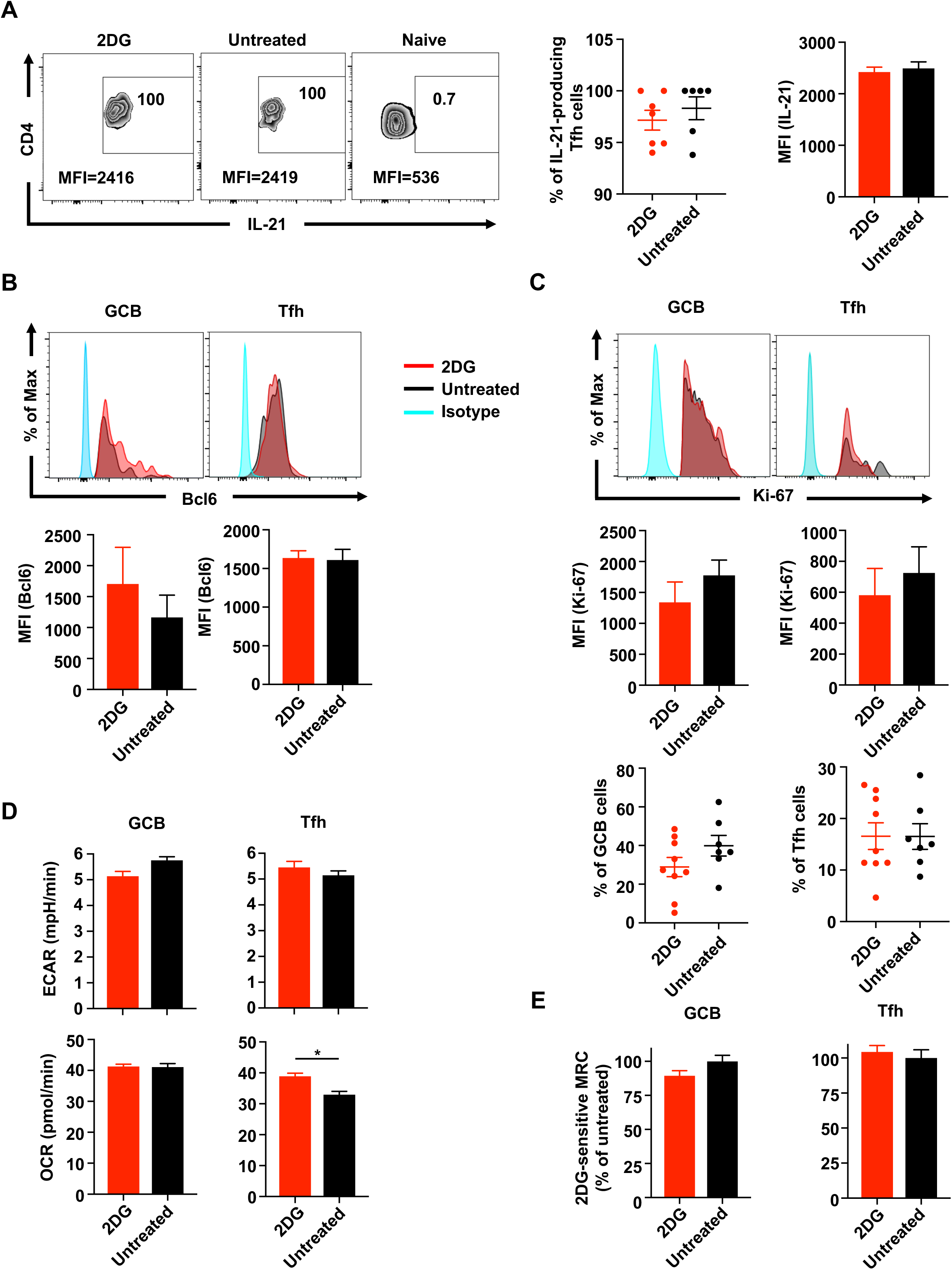
Functionality of autoreactive Tfh and GCB are unaffected by short-term glycolytic inhibition. (**A**) Flow of frequencies and expression of IL-21 in splenic Tfh from symptomatic *Yaa* DKO mice treated with 1-week 2DG and untreated controls. (n= 6-7 mice per group). (**B**) MFI of Bcl6 expression on autoreactive splenic GCB and Tfh from *Yaa* DKO mice treated with 2DG for 1 week or untreated (n= 8-12 mice per group). (**C**) Ki-67 expression and frequency of KI-67^+^ cells in splenic autoreactive GCB and Tfh of *Yaa* DKO mice treated with 2DG for 1 week or untreated (n= 7-9 mice per group). (**D**) Basal ECAR, OCR and (**E**) 2DG-sensitive MRC in GCB and Tfh from 1-week 2DG-treated and untreated *Yaa* DKO mice. Data are from at least three independent experiments. Each dot represents one mouse. Error bars represent mean±SEM; \**p*<0.05, using unpaired two-tailed Student’s *t*-tests.

### BCMA-expressing B cells exhibit elevated glucose dependency and can be targeted to treat lupus

BCMA is a transmembrane glycoprotein exclusively expressed on activated B and plasma cells ^39^, and is associated with disease activity in human lupus ^5,6^. Numbers of total peripheral B cells expressing BCMA were elevated in symptomatic *Yaa* DKO mice compared to pre-symptomatic or healthy mice (**Figure 7A**). Importantly, the frequencies and numbers of peripheral BCMA^+^ GL7^+^ B cells were significantly higher in symptomatic than in pre-symptomatic mice (**Figure 7B**). Following short-term 2DG treatment, numbers of both circulating BCMA^+^ GL7^+^ B cells and splenic BCMA^+^ GCB from symptomatic *Yaa* DKO mice decreased significantly (**Figure 7, C** and **D**). We found that, compared to BCMA^-^ GCB, BCMA^+^ GCB showed significantly higher surface expression of Glut1 (**Figure 7E**), and significantly greater 2-NBDG uptake (**Figure 7F**). Moreover, BCMA^+^ GCB were highly bioenergetic, exhibiting significantly higher basal ECAR and OCR rates compared to BCMA^-^ GCB (**Figure 7, G** and **H**). While both populations showed reduced MRC with 2DG treatment, the reduction was significantly greater in BCMA^+^ GCB than in BCMA^-^ GCB (**Figure 7, I** and **J**). Together, these data demonstrate that BCMA^+^ GCB exhibit a heightened bioenergetic demand and a greater glucose dependency over BCMA^-^ GCB, rendering them particularly susceptible to glycolytic inhibition.

**Figure 7.**
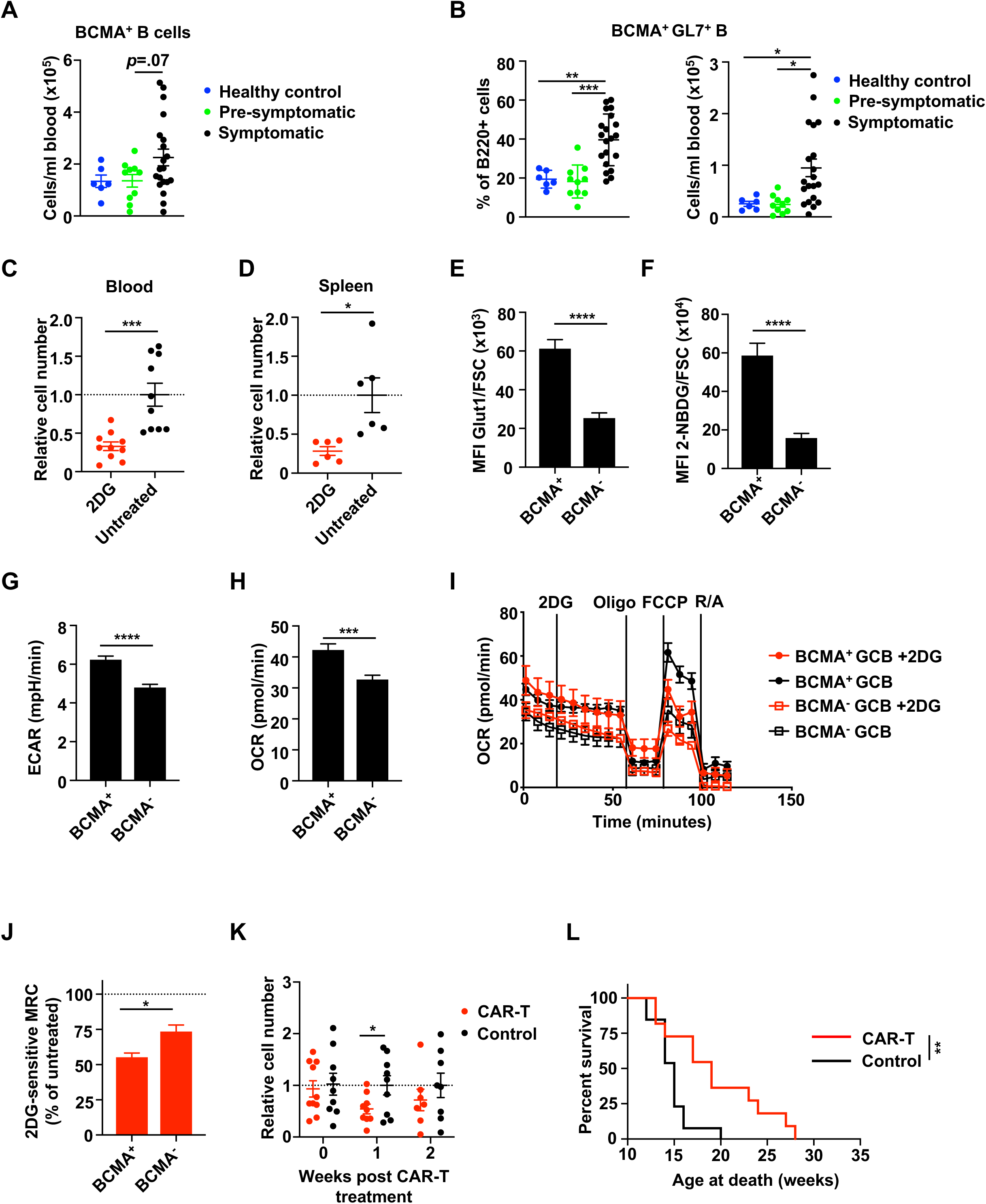
CAR-T cells targeting BCMA reduces GCB and increases lifespan of *Yaa* DKO mice. All experiments were performed with symptomatic *Yaa* DKO mice. Frequencies of BCMA expression on peripheral (**A**) total B cells and (**B**) GL7^+^ B cells from symptomatic, pre-symptomatic and healthy mice (n= 6-20 mice per group). (**C**-**D**) Numbers of (**C**) BCMA^+^ GL7^+^ B cells in blood and (**D**) BCMA^+^ GCB in spleens from mice treated with 2DG for 1 week or untreated, normalized to untreated controls (n=6-10 mice per group). (**E**) Glut1 expression and (**F**) 2-NBDG uptake in splenic BCMA^+^ and BCMA^-^ GCB, normalized to cell size (FSC) (n= 9-10 mice per group). Basal (**G**) ECAR and (**H**) OCR of splenic BCMA^+^ vs. BCMA^-^ GCB. (**I**) OCR and MRC of autoreactive GCB with or without 2DG before treatment with oligomycin (oligo), FCCP, and rotenone/antimycin (R/A). (**J**) Effect of 2DG on MRC as the contribution of glucose metabolism to OXPHOS (2DG-sensitive MRC). (**K**) Relative numbers of peripheral GL7^+^ B cells normalized to untreated controls and (**L**) survival of mice treated with APRIL-based CAR-T or empty vector transduced T cells. Each dot represents one mouse (n= 15 mice per group). Data are presented from at least two independent experiments. Error bars represent mean ± SEM; \**p*<0.05, \*\**p*<0.01, \*\*\**p*<0.001, \*\*\*\**p*<0.0001 using one-way ANOVA with Bonferroni’s multiple comparison tests (**A**-**B, K**), Mantel-Cox test (**I**), or per unpaired two-tailed Student’s *t*-tests (**C**-**H, J**).

To further confirm the role of activated BCMA^+^ B cells in disease, we used chimeric antigen receptor (CAR)-T cells expressing TNFSF13 (APRIL), which is a very high-affinity ligand for BCMA ^39^. By injecting a small number of the APRIL-based CAR-T cells into symptomatic *Yaa* DKO mice with severe clinical disease, we achieved a significant decrease in numbers of circulating activated GL7^+^ B cells after 1 week compared with mice treated with empty vector-transduced T cells (**Figure 7K**). We did not observe decreases in Teff, cTfh or plasmablasts/plasma cell numbers, confirming the specificity of the CAR-T cells (**Supplemental Figure 7**). Mice treated with APRIL-based CAR-T cells exhibited a significantly prolonged mean lifespan (20.4 weeks) compared to control mice (15.6 weeks) (**Figure 7L**). Together, these data support the significant involvement of highly glycolytic BCMA^+^ GCB in lupus pathogenesis and suggest that activated BCMA-expressing B cells can be targeted to treat lupus.

### Glycolysis-related gene expression strongly correlates with the activated B-cell signature, but not the activated T cell signature from lupus patients

To evaluate whether there is elevated glycolysis in the B-cell over T-cell compartment in human SLE, we examined the transcriptomic profile of peripheral CD19^+^ B and CD4^+^ T cells sorted from lupus patients. Linear regression analyses indicated that the gene signature of activated B cells had a strong, positive correlation with glycolysis, significant but weak positive correlations with oxidative phosphorylation, pentose phosphate pathway, tricarboxylic acid cycle and glutamine metabolism, and no correlation with fatty acid oxidation (**Figure 8**). Conversely, the activated T-cell gene signature was only weakly correlated with the pentose phosphate pathway and glutamine metabolism with negative and positive correlations, respectively, but not with glycolysis. When comparing human results to mice, it was noted that peripheral activated B cells from lupus patients and splenic autoreactive B cells of lupus prone mice (**Figure 2B**) were both positively correlated with glycolysis, suggesting that short-term glycolytic blockade may also be efficacious in preferentially targeting pathogenic B cells in human SLE.

**Figure 8.**
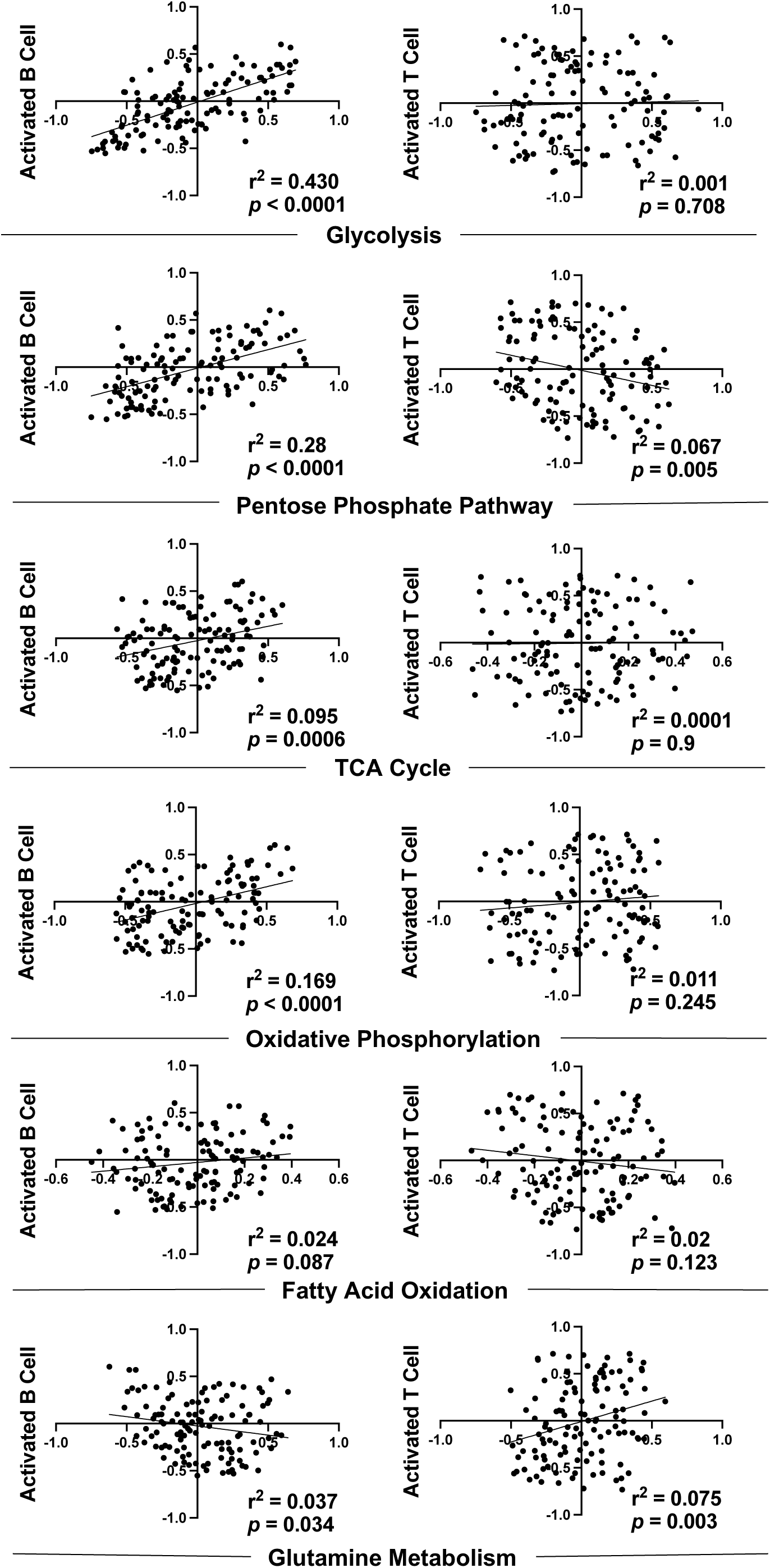
Activated B cells from lupus patients have greater metabolic demand than do activated T cells. Gene-expression data from CD19^+^ B cells or CD4^+^ T cells isolated from lupus patients (n=120 individuals) was used for linear regression between GSVA scores for activated B- or T-cell and metabolic pathway gene signatures. The goodness of fit for each comparison is displayed as the r^2^ value and the slope of the regression line is displayed as the p-value. Correlations with p<0.05 were considered significant. TCA, tricarboxylic acid.

## Discussion

Alterations in glucose metabolism within activated immune/inflammatory cells are increasingly appreciated to underlie lupus and related autoimmune disorders ^40–42^, but how such metabolic modifications are differentially manifested by the various cell types that contribute to lupus pathogenesis remains obscure. Here, we profiled key autoimmune populations in spontaneous lupus models and their response to glycolytic inhibition. We demonstrated a greater glycolytic dependency of GCB compared to that of Tfh. Gene expression signatures for activated B cells were significantly reduced in lupus-prone mice treated with 2DG, whereas signatures for activated T cells remained unaffected. Moreover, glycolysis pathway-related genes were highly correlated with the murine activated B cell signature, but this was weaker with the activated T cell signature. A similar correlation with the glycolysis gene signature was also observed in peripheral activated B cells from human patients with lupus. We found that murine autoreactive B cells, closely resembling GCB, were highly glycolytic and strongly dependent on glycolysis for their survival, but that Tfh were far more metabolically flexible for their energetic requirements. This glycolytic dependency translated into an exploitable weakness whereby pathogenic GCB were preferentially targetable via short-term 2DG treatment. This depletion in lupus-prone mice with advanced clinical disease led to significantly ameliorated renal damage and increased lifespan. Furthermore, we interrogated this lineage in detail, demonstrating that highly activated, BCMA-expressing GCB exhibit heightened glucose metabolism and are exquisitely vulnerable to short-term glycolytic inhibition. Targeting these cells with BCMA-specific CAR-T cells induced lifespan extension equal to that seen in mice treated with 2DG. These findings suggest that glycolytic requirements between autoreactive GCB and Tfh differ and that pathogenic GCB can be selective targets of anti-glycolytic therapy, providing a novel metabolic niche for lupus treatment.

GC formation requires sustained T cell-B cell interaction ^1,9^ and has long been associated with numerous autoimmune conditions ^2^. Long-term treatment of pre-symptomatic *Yaa* DKO lupus-prone mice with 2DG resulted in attenuated cellular disease phenotypes in autoreactive T cells, including Tfh, and markedly reduced GCB and plasma cells/plasmablasts. Moreover, 2DG conferred complete survival for the duration of the treatment in this acute model of lupus. It is evident that long-term glycolytic inhibition via 2DG has a non-specific suppressive effect on a number of cell types, increasing the risk of adverse effects and potentially limiting clinical translatability. Moreover, the broad effects resulting from persistent 2DG treatment *in vivo* make it difficult to identify a key pathogenic cell type that is primarily responsible for reduced lupus manifestations induced by glycolytic inhibition. To elucidate this cell population, we analyzed the metabolic states of distinct autoreactive T and B cells. Our data show that GCB display heightened glycolytic activity (ECAR) over Tfh. Concordantly, GCB exhibited higher Glut1 expression and a correspondingly-elevated glucose uptake compared to Tfh. Elevated Ki-67 expression in GCB over Tfh indicates that these GCB are highly proliferative, which further implicates higher glycolytic rates ^13–15^, directly supporting previous inferences ^41–43^.

Current standard treatment for lupus is non-specific immunosuppression ^40,44^, leaving patients with increased risk of infections and cancers. Preventative effect of long-term 2DG treatment on the activation of numerous autoimmune cell types was previously reported in multiple chronic lupus-prone mice ^16,17^. Recent efforts have focused on targeted therapies for lupus to improve efficacy with fewer adverse effects. For example, belimumab neutralizes the B-cell survival factor TNFSF13B, thereby reducing overall B cell numbers and providing clinical benefit in human lupus ^7^. Here we demonstrate that a 1-week short-term 2DG treatment selectively controls pathogenic GCB in clinically diseased *Yaa* DKO and NZBWF1 mice. Moreover, this GCB reduction served to reverse lupus disease pathology and significantly extend the lifespan of treated acute and chronic models of lupus. Our study also provides the mechanistic explanation by which 2DG reversed abnormal immunophenotypes *in vivo* by demonstrating that glycolytic inhibition caused deficiency of ATP production in pathogenic GCB and a subsequent increase in mitochondrial ROS-mediated metabolic oxidative stress resulting in apoptosis. The increased glycolytic rate in autoreactive GCB may be an adaptation to compensate for excessive ROS production ^45^. Treatment with 2DG impairs mitochondrial function in autoreactive GCB that lack the metabolic flexibility to oxidize other fuels in compensation for glycolytic blockade. Conversely, Tfh exhibit a greater degree of metabolic flexibility to compensate for fuel oxidation during glycolytic restriction, which is likely the reason that they are able to maintain ATP production, survive, and have unimpeded functionality. Furthermore, the lack of 2DG-sensitivity in GCB taken from mice treated with transient 2DG points to the targeted reduction of the pathogenic GCB populations with higher glycolytic dependency, sparing those with lesser glycolytic demand. The evidence of prolonged lifespan of lupus-prone mice with this short treatment 2DG course indicates that glycolysis-dependent autoreactive GCB are pathogenic and 2DG can target these pathogenic GCB populations, thereby avoiding a pan reduction of GCB and other immune cells that would leave the immune system compromised. In addition, the lack of a strong effect on the activated T-cell gene signature in long-term 2DG-treated mice begs the question of the primary mechanism of this glycolytic restriction; especially in view of the interdependency of Tfh and GCB ^2,9^. This presents the possibility that observed Tfh number reductions stem, not from a direct effect of 2DG on Tfh, but on a lack of costimulatory signaling from autoreactive GCB after their depletion via 2DG.

Our findings that autoimmune GCB rely heavily on glucose metabolism differ from those in an immunization paradigm, in which GCB isolated from NP-CGG immunized mice exhibit very low rates of glycolysis and instead rely mainly on FAO to meet their energetic demands ^29^. We observed higher OXPHOS rates in autoreactive GCB over Tfh; however, GCB from lupus mice displayed significantly increased glucose uptake, paired with elevated rates of glycolysis. One reason behind this discrepancy might be that chronic autoantigen stimulation in lupus could cause immune cells to become increasingly activated as disease progresses, versus the more acute stimulation resulting from exposure to a foreign antigen. Moreover, a previous study demonstrated that autoimmune mice treated with 2DG for 9 weeks, and immunized with NP-KLH, maintained a robust GC response against the exogenous antigen, equal to that of untreated, immunized mice, while still exhibiting a reduction in autoreactive GCB numbers ^17^. This suggests that the response of GCB to foreign antigens is refractory to conditions that induce apoptosis in autoreactive GCB, indicating a major disparity in energy requirements between the two GCB types. Further studies are needed to understand this metabolic divergence between autoimmune- and immunization-induced GCB.

Our data has shown highly activated GCB (GL7^+^ B cells) to be the central driver of disease pathogenesis as well as the prime target of anti-glycolytic therapy. Therefore, we further asked the question of what specific cell subtype within this lineage is the main target of 2DG, and found that the subset of BCMA-expressing GCB in lupus-prone mice displayed an extremely high reliance on glycolysis and sensitivity to 2DG. It has been reported that BCMA is overexpressed on both human and murine circulating B cells during lupus progression ^5,46^. Ligation of BCMA is involved in increased expression of the anti-apoptotic gene, *Mcl-1,* in B cells committed to differentiation into long-lived plasma cells ^47,48^. This sustained survival and resistance to apoptosis are likely to be detrimental in lupus. Here, we found elevated BCMA expression on GCB in *Yaa* DKO mice, which was reduced after short-term 2DG treatment, was associated with elevated Glut1 expression and glucose uptake and elevated glycolytic requirements. Indeed, glycolytic inhibition has been reported to decrease Mcl-1 synthesis via promoting the AMPK-mTOR pathway ^49^. Understanding regulation of Mcl-1 levels by glucose metabolism may be a critical aspect of appreciating how glycolytic blockade can differentially affect autoreactive GCB survival. Given their augmented glycolysis usage and association with lupus, BCMA-expressing GCB/GL7^+^ B cells might serve as a sensitive indicator for the efficacy of glycolytic blockade therapy. CAR-T therapy has been used to successfully target refractory, multiple myeloma that highly express BCMA ^39^. Reduced numbers of BCMA-expressing GL7^+^ B cells following our APRIL-based CAR-T cell treatment yielded enhanced survival in treated mice similar to glycolytic inhibition treatment via 2DG, further validating the importance of these cells in lupus pathology. Paired with increase frequencies of BCMA^+^ B cells in lupus patients ^5,6^, hyperactivated BCMA-expressing GCB may be the central B cell population involved in lupus pathogenesis.

In the future, it would be beneficial to interrogate the effect of 2DG treatment in multiple tissues in lupus-prone mice during varied disease stages to understand tissue-specific effects, mechanisms through which lupus develops, and how 2DG ameliorates disease progression in affected tissues. We have not yet investigated whether GCB reduction is sufficient to prevent flares or whether multiple intermittent short-term treatment iterations are required. Although the effects of 2DG on glycolysis are clear, we did not examine to what extent other effects of 2DG, such as altered glycosylation or pentose phosphate pathway, might contribute to the selective reduction of GCB. Finally, murine lupus models have limitations. Our mice critically require the transgenic duplicated expression of *Yaa* (including *TLR7*) or type-I interferon signals. Although these signals are found in 60-80% adult lupus patients ^3^, it is difficult to define how well the findings from available models translate to humans. Nevertheless, glycolytic inhibition with 2DG has been used experimentally in human cancer and COVID-19 clinical trials with surprisingly few significant side effects ^50–52^. Since glycolysis is highly conserved between mice and humans, and we found similar gene expression profiles in activated human and murine lupus B cells, the chances of a successful therapeutic translation of 2DG treatment to lupus patients are substantial.

Overall, this work illustrates that autoreactive GCB have a greater glycolytic rate and dependency, and pathogenic GCB retain apoptotic sensitivity to glycolysis inhibition compared with Tfh. Because of this differential requirement, we demonstrated that a 1-week course of 2DG specifically targets BCMA-expressing GCB with high glycolytic dependency, and that the reduction of this subset confers significant amelioration of lupus pathology, significantly extending survival, all while preserving the number and function of other immune cells. Future treatments designed to target these pathogenic GCB, including short-term glycolytic inhibition, may improve the outcomes of patients suffering from lupus and other autoimmune disorders.

## METHODS

### Mice and treatments

BXSB.Cg-*Cd8a^tm1Mak^ Il15^tm1Imx^*/Dcr *Yaa* (*Yaa* DKO), NZBWF1/J (JAX#100008), BXSB.B6-*Yaa*^+^/MobJ (JAX#001925), and C57BL/6J (JAX#000664) mice were bred and housed at The Jackson Laboratory (Bar Harbor, Maine). Mice were provided 10% fat food JL Mouse Breeder/Auto (LabDiet**^®^** 5K20) and water *ad libitum*; they were housed on a 14-hour light, 10-hour dark cycle in a specific pathogen-free room. Symptomatic male *Yaa* DKO mice had visibly swollen lymph nodes and displayed elevated cellular disease biomarkers (often found at age 10-12 weeks old); whereas pre-symptomatic males (6-7 weeks old) had yet to develop these manifestations. Female littermate mice lacking the *Yaa*-containing Y chromosome are immunologically normal and were used as healthy controls ^18^. Before experiments, mice were bled to verify disease state. Symptomatic NZBWF1 mice, were 36-40-week-old females with elevated protein urea. Experiments were performed on symptomatic mice unless otherwise stated. Mice in survival experiments were euthanized when classified as moribund, and mice euthanized before this point or for non-lupus-related issues were excluded from survival data. For *in vivo* metabolic treatments, 2DG (Thermo Fisher, 6 g/L) was dissolved in drinking water and was provided to mice *ad libitum*. Animal studies were approved by Institutional Animal Care and Use Committee at The Jackson Laboratory.

### Flow cytometry and intracellular staining

Single-cell suspensions from peripheral blood, spleens, or lymph nodes were blocked with anti-CD16/32 and stained with antibody cocktails on ice after red blood cell lysis. Flow data was acquired on an LSR II flow cytometer and was analyzed using FlowJo software version 10.6.1. Intracellular staining was performed with the Fixation/Permeabilization Solution Kit (BD Biosciences) using manufacturer’s recommendations with cells stained overnight on ice. Cell populations were either used directly or further purified by a FACSAria II cell sorter. For apoptosis detection, the FITC-Annexin V/propidium iodide (PI) double staining assay was used. Cells were stained for surface markers as described, followed by 2x washing with Annexin staining buffer (PBS with 10 mM HEPES, 150 mM NaCl and 5 mM CaCl_2_). Cells were then stained at room temperature for 10 minutes in the dark before flow analysis. For measuring mitochondrial mass, mitochondrial membrane potential, ROS and ATP levels, cells were stained with MitoTracker Green FM, MitoTracker Red, MitoSOX (Life Technologies), and BioTracker ATP-Red (MilliporeSigma), respectively, according to the manufacturer’s instructions.

### Cell survival assays

*Yaa* DKO splenocytes or healthy human PBMCs (Precision for Medicine, MD) were cultured at 4×10^6^ per ml of R10 medium (RPMI plus 10% FBS, 2 mM L-glutamine, 50 μM 2-ME, 100 U/ml penicillin, and 100 μg/ml streptomycin) with 0.5 μg/ml of anti-CD3/28 antibodies, 100 U/ml IL-2 and 30 ng/ml of the TLR7 agonist R848 (Adipogen) for 2 days to achieve cellular activation ^53^. On day 2, the cells were harvested and plated at 2.5×10^6^ cells per ml in R10 medium with 0, 0.2, or 1 mM 2DG onto a 96-well plate at 200 μl per well. Cells were stained with antibody cocktail and analyzed by flow cytometry to ascertain numbers of live cells of each type by PI staining. For measuring viability of autoreactive cells, indicated cells were sorted from the splenocytes of symptomatic *Yaa* DKO mice and were plated at 1×10^5^ cells per ml in R10 media with or without indicated glycolysis inhibitors for 1-2 days. Cells were harvested and analyzed by flow cytometry to ascertain live cell numbers.

### Metabolic assays

Oxygen consumption rates (OCR) and extracellular acidification rates (ECAR) were measured in Seahorse XF assay media (non-buffered RPMI 1640 containing 25 mM glucose, 2 mM L-glutamine, and 1 mM sodium pyruvate) under basal conditions and in response to 25 mM 2DG, 20 μM UK5099, 10 μM BPTES, 40 μM etomoxir, or 100 nM thioridazine with the addition of 1 μM oligomycin, 1.5 μM fluoro-carbonyl cyanide phenylhydrazone (FCCP), and 100 nM rotenone plus 1 μM antimycin A (Sigma) with a XF-96^e^ Extracellular Flux Analyzer (Seahorse Bioscience). Cellular rates of ATP production were measured by Seahorse Real-Time ATP Rate kit and calculated according to manufacturer’s recommendations. All assays were performed with equal numbers of cells per well in each group. Maximal respiratory capacity (MRC) was assessed as the percentage of baseline OCR after FCCP injection, and the maximum glycolytic capacity (MGC) was calculated as the percentage of baseline ECAR after oligomycin injection. For analysis of glucose uptake, mice were injected with 50 μg of 2-NBDG via tail vein 20 minutes before spleens were harvested. Cells were stained for surface markers and intensity of 2-NBDG in each population was assessed by flow cytometry. Glucose concentrations were conducted using a Glucose Assay Kit from Eton Biosciences according to manufacturer’s specifications and readings were carried out using a SpectraMax i3 spectrophotometer (Molecular Devices) at 490nm absorbance.

### Immunohistochemistry

Six-micron sections were cut from fresh spleen tissue frozen in OCT medium and were mounted on slides, fixed with ice-cold acetone for 15 minutes, and re-hydrated for 10 minutes with PBS. Leica SuperBlock was used to block the slides for 30 minutes before staining with PNA at 20 μg/ml and IgD at 5 μg/ml for 1 hour at room temperature in the dark. Slides were washed 3 times with PBS before mounting sections with Prolong™ Diamond Antifade Mountant (Invitrogen) and were imaged within 24 hours of staining on a Leica Thunder imager per manufacturer’s recommendations. Individual images were merged using the ‘mosaic merge’ function in Leica Application Suite X software and were exported into Fiji for image analysis. GCs were identified as areas of PNA staining surrounded by IgD-stained cells. GC size was calculated by measuring the area of PNA stain within each GC three times and taking the average of these replicates for the final area.

### Gene expression analysis by RNA-sequencing and quantitative real-time PCR

RNA-sequencing (RNA-seq) was performed by The Jackson Laboratory Genome Technologies Scientific Service. Total RNA of spleens (4 samples per group) was isolated with the QIAGEN miRNeasy mini extraction kit (QIAGEN) and cDNA was synthesized with the High Capacity cDNA Reverse Transcription Kit (Applied Biosystems). RNA quality was assessed with a Bioanalyzer 2100 (Agilent Technologies). Poly(A)-selected RNA-seq libraries were generated using the Illumina TruSeq RNA Sample preparation kit v2. RNA-seq was performed in a 75-bp paired-end format with a minimum of 40 million reads per sample on the Illumina NextSeq 500 platform according to the manufacturer’s instructions. RNA-seq reads were filtered and trimmed for quality scores >30 using a custom python script. The filtered reads were aligned to *Mus musculus* GRCm38 using RSEM (v1.2.12) ^54^ with Bowtie2 (v2.2.0) ^55^ (command: rsem-calculate-expression -p 12 --phred33-quals --seed-length 25 --forward-prob 0 --time --output-genome-bam -- bowtie2). RSEM calculates expected counts and transcript per million (TPM). The expected counts from RSEM were used in the Bioconductor edgeR 3.20.9 package ^56^ to determine differentially expressed genes. Determination of the most significantly different gene groups was performed with Gene Set Enrichment Analysis (GSEA) ^24,25^. The following gene sets from the Molecular Signatures Database were used: systematic names, M12175 (Systemic Lupus Erythematosus) and M12930 (Immunoglobin Production). Groups were deemed significant at an FDR *q*-value of 0.05 or below. Genes of interest for each group were extracted based on the core enrichment value provided by GSEA. Quantitative real-time PCR (qPCR) was performed by the SYBR green method, with an Applied Biosystems ViiA 7 Real-Time PCR System (Life Technologies). Quantitation of the results was performed by the comparative Ct (2-[delta][delta]Ct) method. The Ct value for each sample was normalized by the value for beta-actin. The sequences of qPCR primers are listed in **Supplemental Table 1**.

### Gene set variation analysis (GSVA)

The R/Bioconductor software package GSVA ^26^ (v1.25.0) was used as a non-parametric, unsupervised method to estimate enrichment of pre-defined gene sets in RNA-seq data from *Yaa* DKO mice. The inputs for GSVA were a matrix of log2 gene expression values for all samples and curated gene sets describing select immune cell types and pathways. Low-intensity probes were filtered out if the interquartile range (IQR) of their log2 expression values across all samples was not greater than zero. Enrichment scores were calculated using a Kolgomorov Smirnoff (KS)-like random walk statistic and represented the greatest deviation from zero for a particular sample and gene set. Scores across all samples were normalized to values between -1 (indicating no enrichment) and +1 (indicating enrichment). Significant differences in enrichment between 6wk, 10wk, and 10wk+2DG cohorts were calculated using Brown-Forsythe and Welch ANOVA tests with Dunnett T3 test for multiple comparisons with an alpha of 0.05.

### GSVA gene set generation and co-expression analysis

Mouse gene sets used as input for GSVA are listed in **Supplemental Tables 2 and 3**. Immune-cell and functional-pathway gene signatures were generated by an iterative process involving derivation through literature mining, gene ontology (GO) term definitions provided by the Mouse Genome Informatics (MGI) GO Browser ^57^, and immune-cell expression as determined by the Immunological Genome Project Consortium (ImmGen) ^58^ followed by dataset-specific curation by co-expression analysis. The following GO terms were used to generate the Activated B Cell, Pentose Phosphate Pathway, and Glutamine Metabolism gene sets: GO:0002312-B cell activation involved in immune response, GO:0006098-Pentose phosphate shunt, and GO:0006541 glutamine metabolic process. The glycolysis, oxidative phosphorylation, TCA cycle, and fatty acid oxidation gene signatures have been previously described ^59^. The B-cell, plasma cell, T-cell, activated T-cell, and myeloid-cell signatures were derived from Mouse CellScan, a tool for the identification of cellular origin of mouse gene-expression datasets. For human data, gene expression analysis was performed on publicly available transcriptomic data (GEO accession: GSE164457) ^60^ from participants recruited from the California Lupus Epidemiology Study. Briefly, PBMCs were isolated from 120 lupus patients, and were sorted into populations of CD19^+^ B cells and CD4^+^ T cells for bulk RNA-seq. GSVA was performed using human orthologs of mouse gene sets listed in **Supplemental Tables 4 and 5**. Linear regression analysis between GSVA enrichment scores for activated B-cell, activated T-cell, and metabolic pathway gene signatures was carried out as previously described for the mouse RNA-seq data. Co-expression analysis for all gene sets was carried out in R using the cor() function to compute Pearson correlation coefficients for log2 expression values of each gene in a given gene set. Gene sets were then refined to maximize the number of co-expressed genes while maintaining specificity for a given cell type or functional pathway. Linear regression analysis between GSVA enrichment scores for immune cell and functional pathway gene signatures was carried out using GraphPad Prism software (v9.0.1). For each analysis, the goodness of fit is displayed as the R^2^ value. The significance of the slope of the regression line is displayed as the *p*-value.

### Retrovirus vector designs and generation of APRIL-based CAR-T cells

The cDNA sequence of the extracellular region of mouse APRIL (nucleotides 319-721, GenBank Q9D777) was fused to the transmembrane/intracellular domain of CD28 (nucleotides 543–740, GenBank NM_007642.4) and intracellular domain of CD3ζ (nucleotides 302–643, GenBank NM_001113391.2) via the hinge of mouse IgG1 and custom-synthesized by Integrated DNA Technologies. The synthesized cDNA was cloned in-frame into the MSCV-IRES-GFP-based retroviral vector ^14^. For generating retroviruses, APRIL or empty control vectors were transfected into 293T cells (together with helper plasmid) and the resulting culture viral supernatants were used to transduce CD8^+^ T cells isolated from BXSB.B6-*Yaa*^+^ mice, a BXSB-consomic strain that does not carry the *Yaa* mutation and does not develop lupus-like disease. Cells were activated with anti-CD3/28 antibodies for 1 day and exposed to viral supernatant for 90 minutes (at 2,500 rpm, 30°C) in medium containing hexadimethrine bromide (8 μg/mL), HEPES (1 mM) and IL-2 (100 U/mL). Cells were cultured for an additional 2 days before sorting for GFP-expressing cells, which was the marker for retroviral expression.

### Adoptive transfers

For adoptive transfer of autoreactive GCB, symptomatic *Yaa* DKO mice were pre-treated with 2DG for 1 week and were then removed from treatment. Two days after removal, each mouse was injected intravenously with 1.5×10^6^ GCB or non-GCB cells sorted from untreated, symptomatic *Yaa* DKO mice. For adoptive transfer of APRIL-based CAR-T cells, 1×10^6^ transduced T cells were injected intravenously into symptomatic *Yaa* DKO mice.

### Statistical analysis

Comparisons for two groups were calculated by using an unpaired, two-tailed Student’s *t*-test. Comparisons for more than two groups were calculated using a one-way ANOVA followed by Bonferroni’s multiple comparison tests. Kaplan-Meier curve with Log-rank (Mantel-Cox) test was used for comparisons of survival curves. GSVA analysis comparisons were calculated using Brown Forsythe and Welch’s ANOVA followed by Dunnett’s T3 multiple comparisons test. All statistical tests were conducted in GraphPad Prism V8.0.2.

## ACKNOWLEDGMENTS

We thank Will Schott, Danielle Littlefield, and Krystal-Leigh Brown for procedural expertise with flow cytometry experiments, Philipp Henrich for technical assistance with fluorescent imaging, Karolina Palucka, Guangwen Ren and David Serreze for reagents and advice, and Edison Liu for critical review, and Stephen Sampson for critical editing of the manuscript. This work was funded by The Jackson Laboratory Director’s Innovation Fund (19000-18-19) to CC, and supported by the RILITE Foundation, and the John and Marcia Goldman Foundation to PEL.

## AUTHOR CONTRIBUTIONS

JJW, JW, LB, ACG., PEL, DCR and CC designed the research. JJW, JW, JDS, EB, CLM and CHC performed experiments and analyzed data. LB, ACG and CC gave conceptual advice. ARD and GAS performed computational analysis. All authors contributed to writing the manuscript.

## MATERIALS & CORRESPONDENCE

Address correspondence to: Chih-Hao Chang, The Jackson Laboratory, Bar Harbor, Maine 04609, USA.

## Methods

### Flow cytometry and intracellular staining

All antibodies were purchased from BioLegend or eBioscience. The following markers were used to identify each *ex vivo* population: B220^+^ CD4^-^ (total B cells), B220^+^ CD4^-^ CD138^-^ GL7^+^ (GCB/activated GL7^+^ B cells), B220^+^ CD4^-^ CD138^-^ GL7^-^ (non-GCB), B220^+^ CD4^-^ CD138^+^ GL7^-^ (plasmablasts/plasma cells), B220^-^ CD4^+^ (total CD4 T cells), B220^-^ CD4^+^ CXCR5^+^ PD1^+^ (Tfh), B220^-^ CD4^+^ CD62L^-^ CD44^+^ (Teff), B220^-^ CD4^+^ CD44^-^ CD62L^+^ (naïve T cells), B220^-^ CD4^+^ CXCR5^+^ PD1^+^ BCL6^+^ Foxp3^+^ (Tfr), B220^-^ CD4^+^ CD25^+^ Foxp3^+^ (Treg), B220^+^ CD4^-^ CD21^-^ CD23^-^ (T1), B220^+^ CD4^-^ CD21^+^ CD23^+^ (T2), B220^+^ CD4^-^ CD21^+^ CD23^-^ (MZ), B220^+^ CD4^-^ CD21^-^ CD23^+^ (FO) and B220^-^ CD4^-^ CD11b^+^ (myeloid cells). The following markers were used to identify each mouse *in vitro* activated cell population: CD4^-^ B220^+^ GL7^+^ (GCB-like cells), B220^-^ CD4^+^ CD62L^-^ CD44^+^ (activated T/Teff-like cells). The following markers were used to identify each human *in vitro* population: CD4^-^ CD20^+^ IgD^-^ BR3^+^ (GCB-like cells) and CD20^-^ CD4^+^ CD45RA^-^ CD45RO^+^ (activated T/Teff-like cells).

---

**Supplemental Figure 1.**
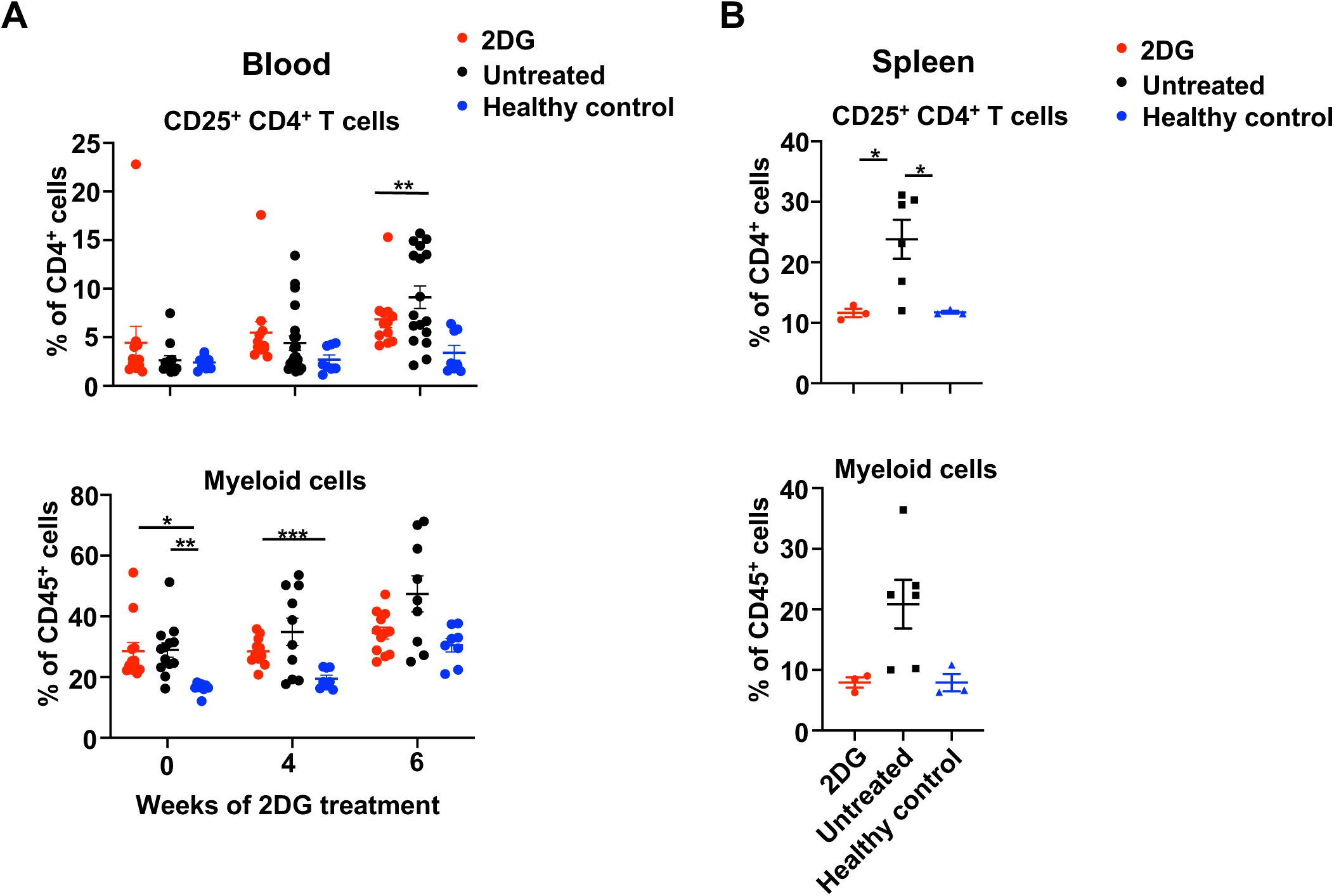
Long-term glycolytic inhibition decreases circulating and splenic cellular populations in lupus-prone mice. (**A**) Flow cytometric results of peripheral CD25^+^ T and myeloid cells of *Yaa* DKO mice treated with 2DG at 6-week-old for indicated weeks or untreated, and untreated healthy controls, respectively (n=15-25 mice per group). (**B**) Flow cytometric results of splenic cells taken from 13-week old *Yaa* DKO mice treated with 2DG for 7 weeks or untreated, and healthy controls (n=3-8 mice per group). Data are representative from at least two independent experiments. Each dot represents one mouse. Error bars represent mean±SEM; *p<0.05, **p<0.01, \*\*\**p*<0.001, \*\*\*\**p*<0.0001 using one-way ANOVA with Bonferroni’s multiple comparison tests.

**Supplemental Figure 2.**
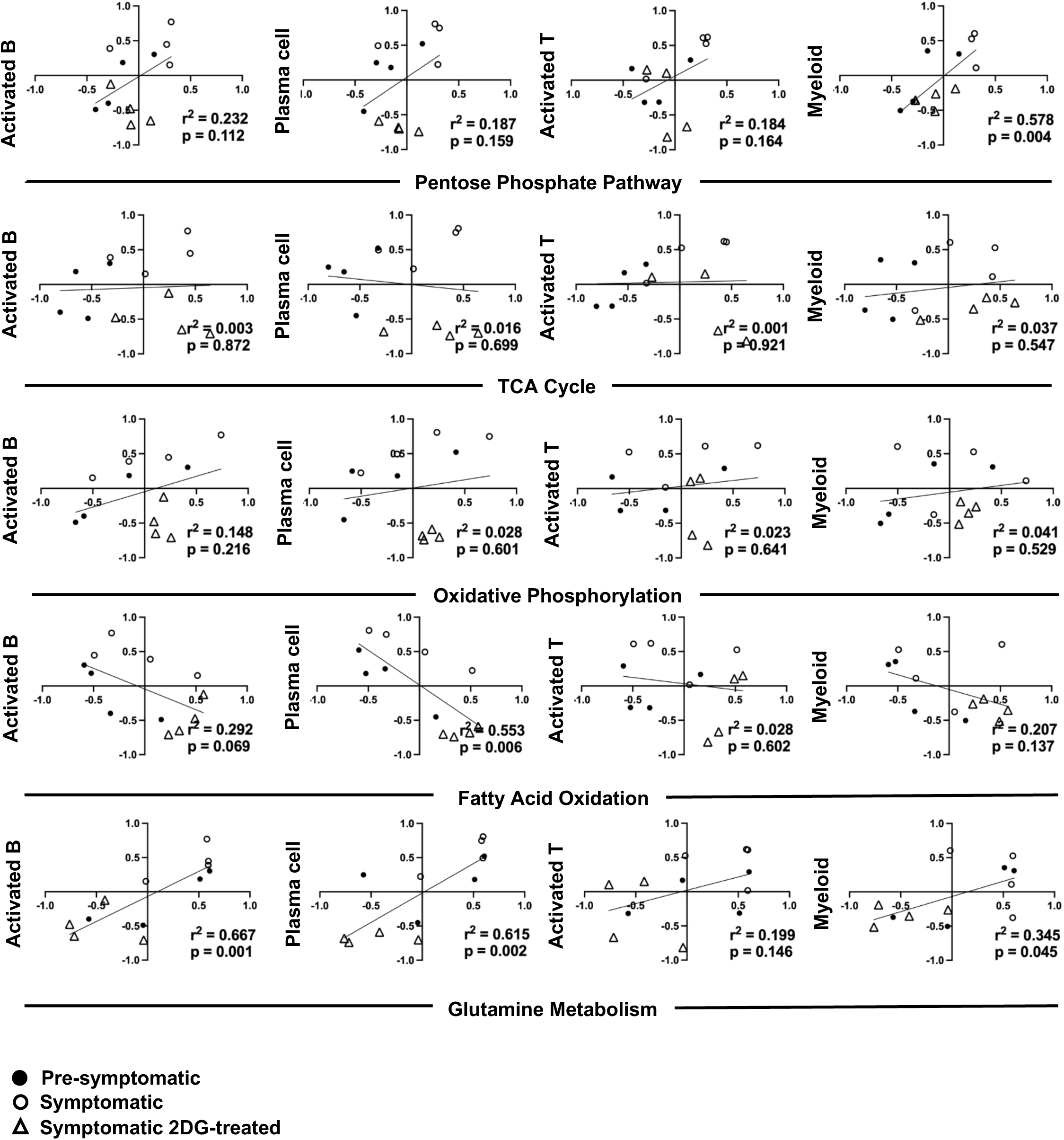
Analysis of the effect of long-term glycolytic inhibition on lupus T/B-cell populations. Linear regression between GSVA scores for activated T/B cells, plasma, and myeloid cells compared to various metabolic pathways. Data are from splenocytes of individual 6-week-old untreated (pre-symptomatic) *Yaa* DKO mice (●), or 10-week-old (Symptomatic 2DG-treated) *Yaa* DKO mice treated with 2DG for 4 weeks (○), or untreated (Symptomatic) (△). (n=4 mice per group). The goodness of fit for each comparison is displayed as the r^2^ value and the slope of the regression line is displayed as the p-value. Correlations with p<0.05 were considered significant. TCA, tricarboxylic acid.

**Supplemental Figure 3.**
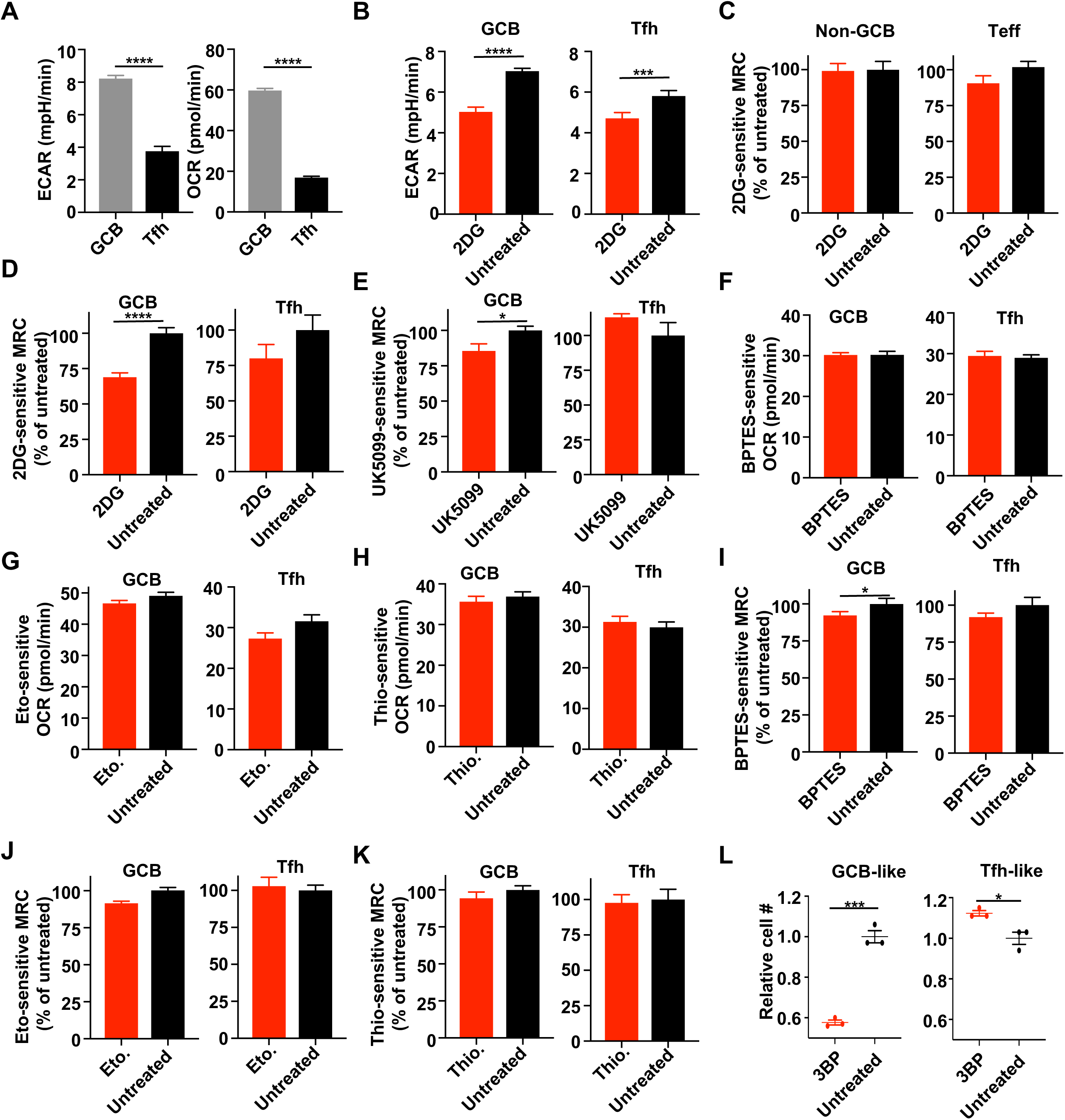
Autoreactive GCB display lower metabolic flexibility than Tfh and are highly dependent on glucose metabolism. Metabolic flux profiling was assessed on splenic GCB and Tfh sorted from either NZBWF1 mice (**A**, **D)** or *Yaa* DKO (**B, C**, **E**-**K**). (**A**) Basal OCR and ECAR of Tfh and GCB. (**B**) Basal ECAR, of Tfh and GCB. The effect of 2DG on MRC (2DG-sensitive MRC) of, (**C**) non-GCB and Teff, and of (**D**) Tfh and GCB. (**E**) The effect of UK5099 on the contribution of glycolysis to OXPHOS (UK5099-sensitive MRC). The effect of BPTES, etomoxir (Eto.) or thioridazine (Thio.) treatment on (**F**-**H**) basal mitochondrial respiration (OCR), or (**I**) the contribution of glutaminolysis (BPTES-sensitive MRC), (**J**-**K)** FAO (Eto-sensitive MRC, or Thio-sensitive MRC). (**L**) Live-cell (PI^-^) numbers of GCB-like and Tfh-like cells after 24-hour treatment with 1 mM 3-bromopyruvate (3BP). Data are from at least three independent samples. Error bars represent mean ± SEM; \**p*<0.05, \*\*\**p*<0.001, \*\*\*\**p*<0.0001 using unpaired two-tailed Student’s *t*-tests.

**Supplemental Figure 4.**
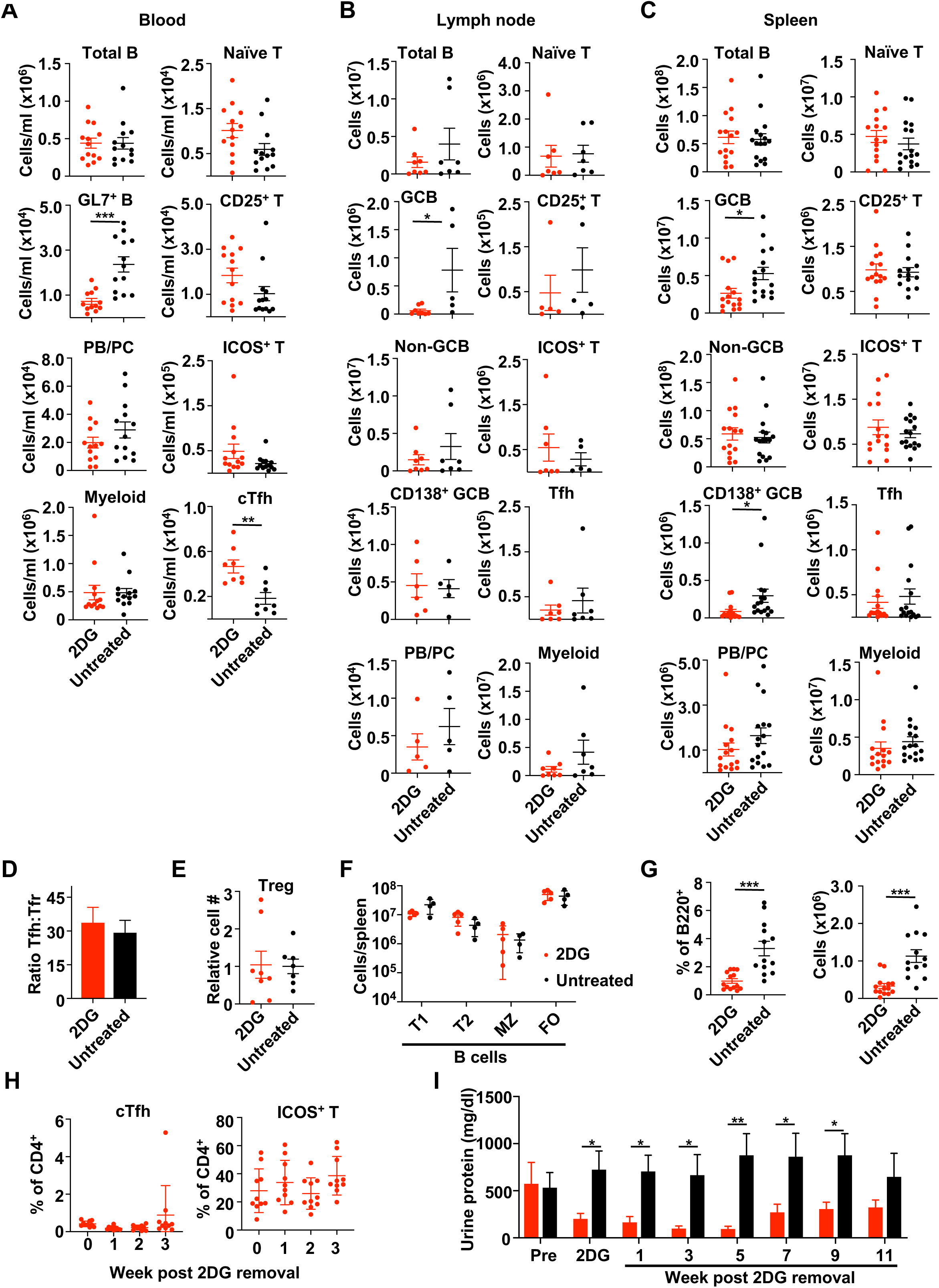
Short-term glycolytic inhibition causes targeted reductions of autoreactive GCB, and decreased urine protein in lupus-prone mice. Flow cytometric data on cell numbers from *Yaa* DKO mice treated with 2DG for 1 week compared to untreated mice in (**A**) blood, (**B**) lymph nodes and (**C**) spleens (n= 5-13). PB/PC, plasmablasts/plasma cells. (**D**) Ratio of splenic Tfh numbers to Tfr-cell numbers (n=4-5). (**E**) Relative numbers of splenic Treg cells (n=7-8). (**F**) Numbers of splenic transitional type 1 (T1), 2 (T2), marginal zone (MZ) or follicular (FO) B cells per spleen (n=4-5). (**G**) Frequencies (left) and numbers (right) of splenic B220^+^ CD95^hi^ CD38^low^ B cells (n=13-14). (**H**) Circulating Tfh (cTfh) and ICOS^+^ CD4^+^ T cells on day 7 of 2DG treatment (week 0) and 1-3 weeks after 2DG removal (n= 10). (**I**) Urine protein concentrations from NZBWF1 mice, in which 1-week of 2DG treatment was started at 36 weeks of age (n= 13-16). Data are from at least three independent samples. Each dot represents one mouse. Error bars represent mean±SEM; \**p*< 0.05, \*\**p*<0.01, \*\*\**p*< 0.001 using unpaired two-tailed Student’s *t*-tests (**A**-**E, G**) or one-way ANOVA with Bonferroni’s multiple comparison tests (**F**, **H**-**I**).

**Supplemental Figure 5.**
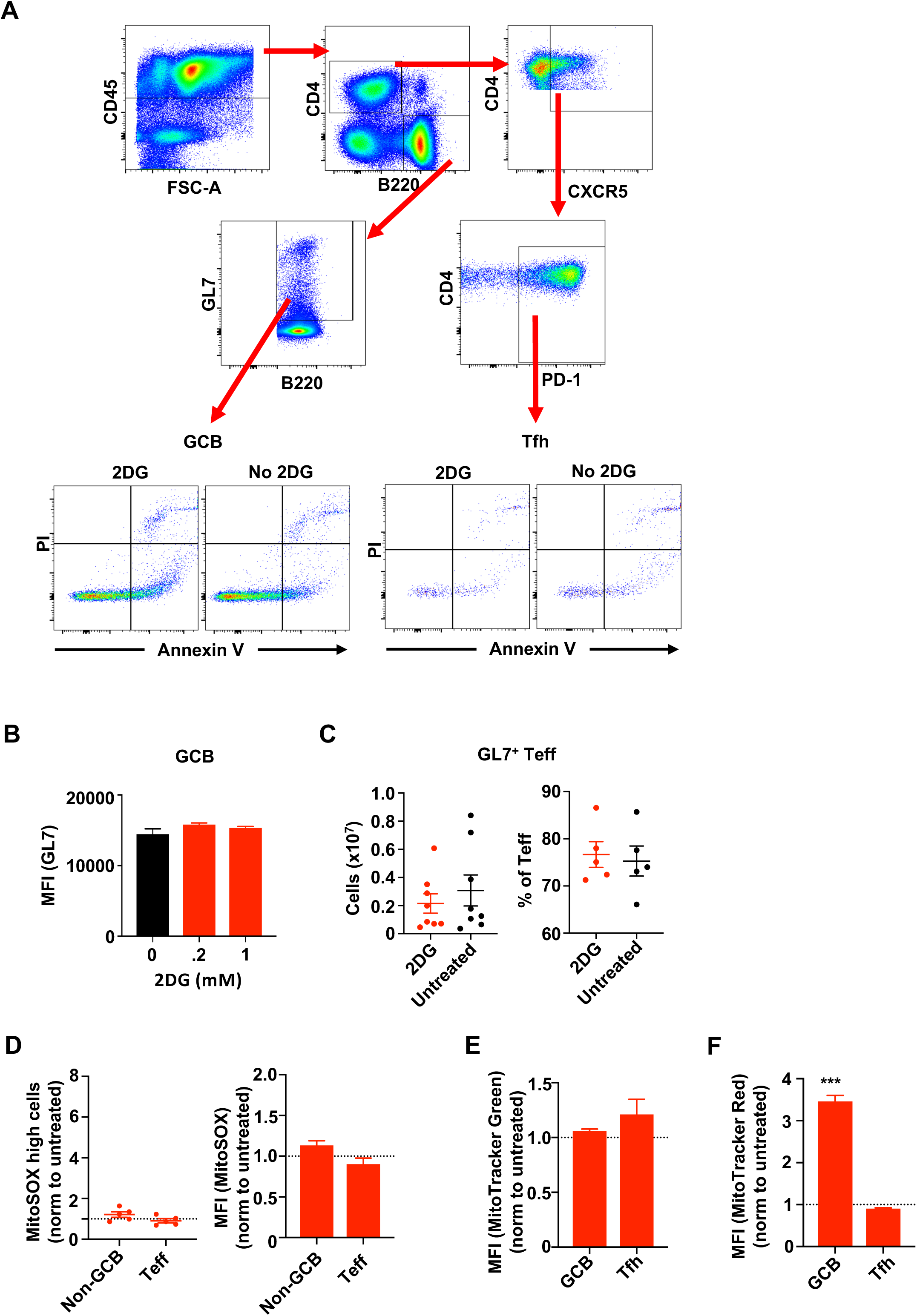
Short-term 2DG exposure does not alter GL7 expression and mitochondrial mass in autoreactive GCB or T cells. (**A**) Representative gating strategy for apoptotic determination via annexin V and propidium iodide (PI) staining. (**B**) Mean fluorescence intensity (MFI) of GL7 expression on splenic GCB treated with 0, 0.2, or 1 mM 2DG for 1 day (n=3). (**C**) Flow data on numbers and frequencies of splenic Teff expressing GL7 in *Yaa* DKO mice treated with 2DG for 1 week compared to untreated controls (n=8). (**D**) Frequencies and MFI of mitochondrial ROS (MitoSOX) in splenic non-GCB and Teff of *Yaa* DKO mice treated with 2DG for 1 week normalized to untreated controls (n=5). (**E**) Mitochondrial mass and (**F**) membrane potential assessment of splenic GCB and Tfh via MitoTracker Green or Red relative to untreated controls (n=5). Data are from at least three independent samples. Error bars represent mean±SEM; \*\*\**p*< 0.001 using one-way ANOVA with Bonferroni’s multiple comparison tests (**B**) or unpaired two-tailed *t*-tests (**C**-**F**).

**Supplemental Figure 6.**
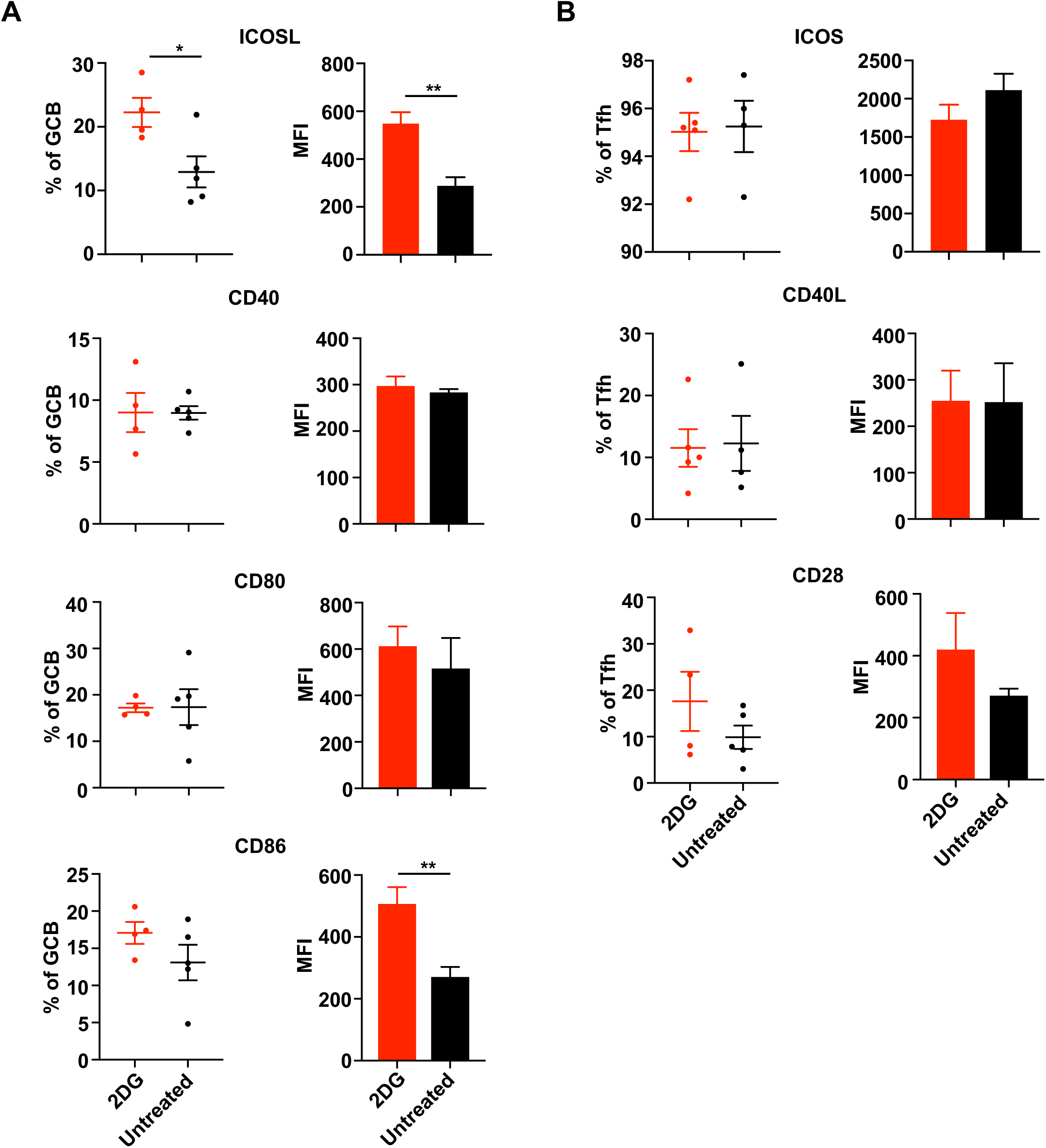
Short-term 2DG treatment does not inhibit Tfh-GCB interaction molecules. Flow analysis of the frequency and MFI of (**A**) ICOS-L, CD40, CD80 and CD86 markers on the surface of GCB and (**B**) ICOS, CD40L and CD28 on the surface of Tfh from *Yaa* DKO mice treated with 2DG for 1 week compared to untreated controls (n= 4-5 mice per group). Each dot represents one mouse. Data are from two independent experiments. \*\**p*<0.01, using unpaired two-tailed Student’s *t*-tests. Error bars represent mean±SEM.

**Supplemental Figure 7.**
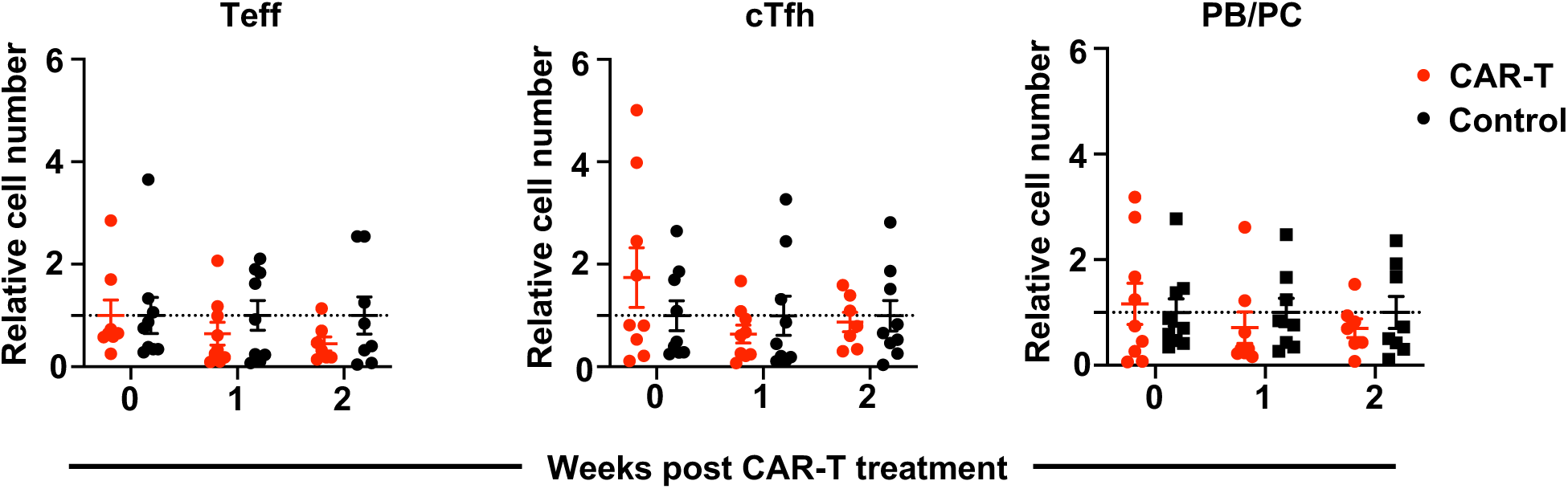
Teff, Tfh, and plasmablasts/plasma cells are unaffected by APRIL-based CAR-T cell therapy. Relative numbers of peripheral Teff, cTfh, and plasmablasts/plasma cells (PB/PC) from *Yaa* DKO mice after treatment with APRIL-based CAR-T cells or control T cells (n= 9 mice per group). Data are from two independent experiments. Each dot represents one mouse. Error bars represent mean±SEM. No statistically significant (*p*>0.05) difference using one-way ANOVA with Bonferroni’s multiple comparison tests.

**Supplemental Table 1.**
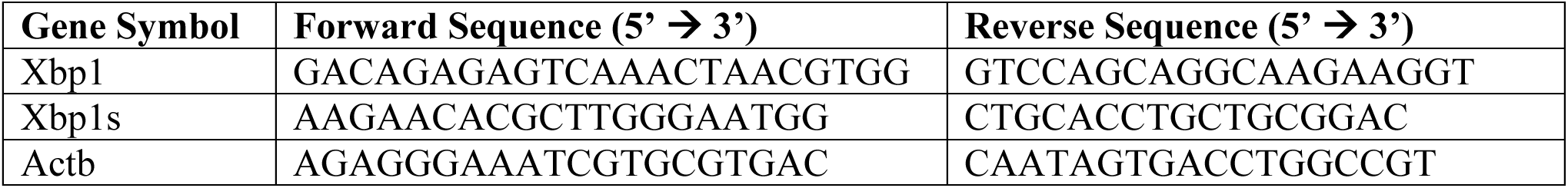
Primer sequences.

**Supplemental Table 2.**
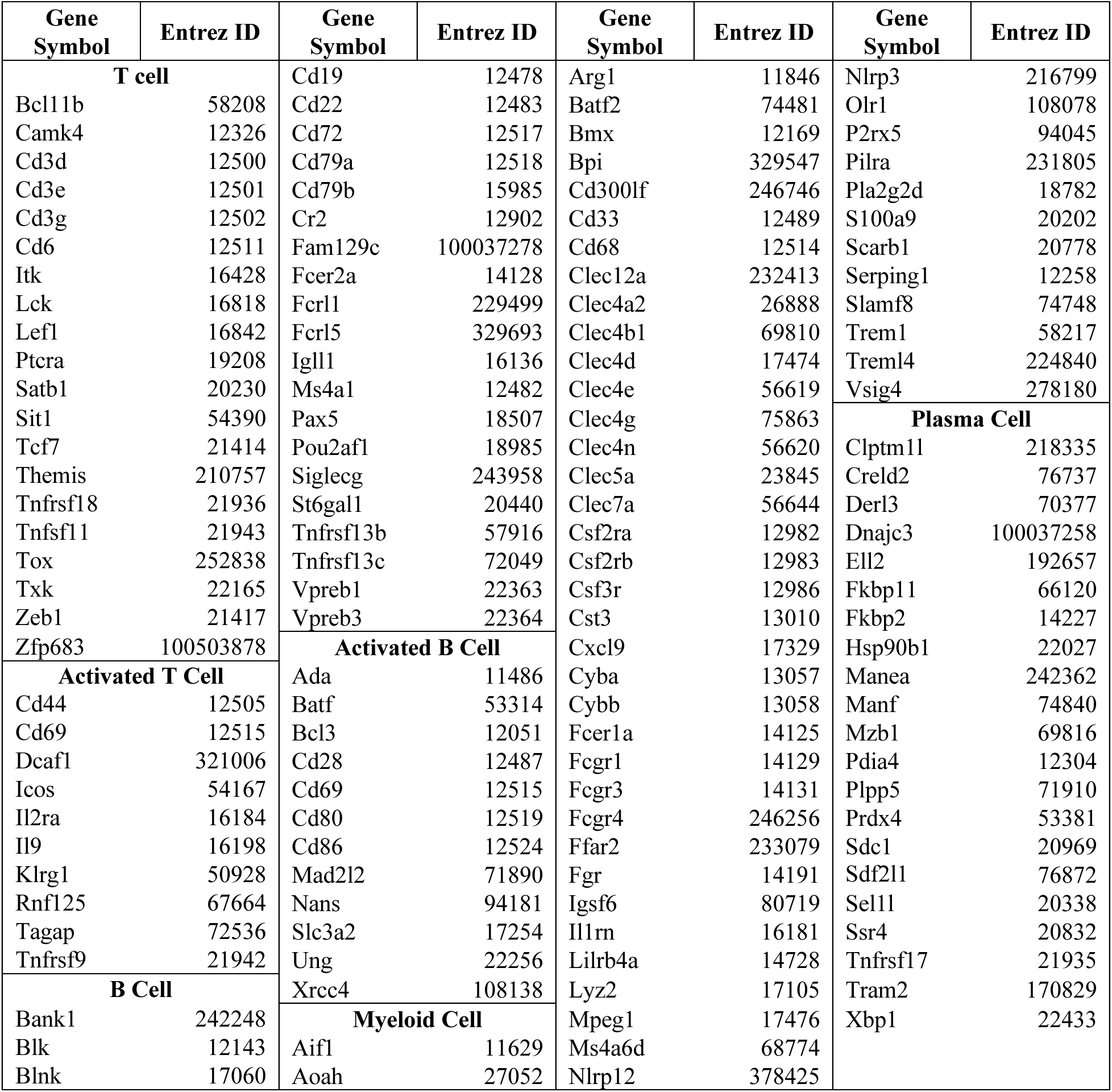
Input list of murine gene sets of immune cells for Gene Set Variation Analysis.

**Supplemental Table 3.**
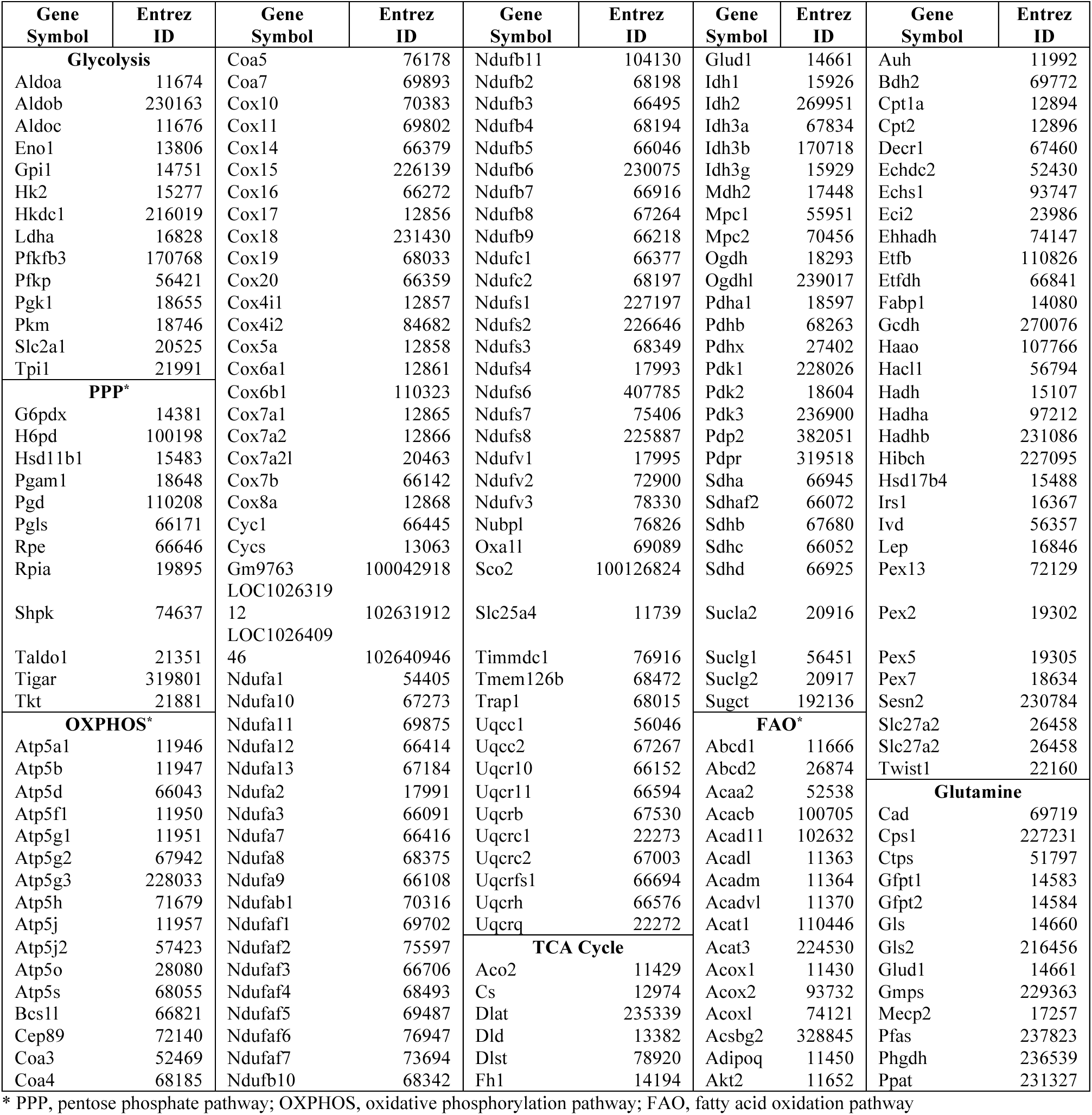
Input list of murine gene sets of metabolic pathways for Gene Set Variation Analysis.

**Supplemental Table 4.**
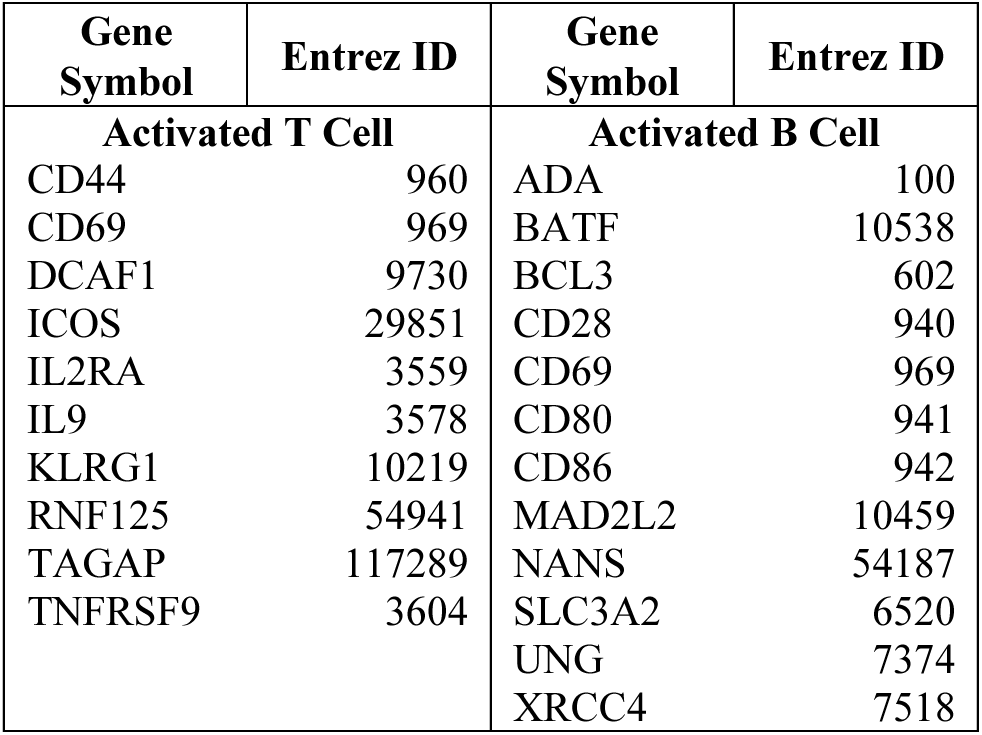
Input list of human gene sets of immune cells for Gene Set Variation Analysis.

| **Gene Symbol** | **Entrez ID** | **Gene Symbol** | **Entrez ID** |
| --- | --- | --- | --- |
| **Activated T Cell** |  | **Activated B Cell** |  |
| CD44 | 960 | ADA | 100 |
| CD69 | 969 | BATF | 10538 |
| DCAF1 | 9730 | BCL3 | 602 |
| ICOS | 29851 | CD28 | 940 |
| IL2RA | 3559 | CD69 | 969 |
| IL9 | 3578 | CD80 | 941 |
| KLRG1 | 10219 | CD86 | 942 |
| RNF125 | 54941 | MAD2L2 | 10459 |
| TAGAP | 117289 | NANS | 54187 |
| TNFRSF9 | 3604 | SLC3A2 | 6520 |
|  |  | UNG | 7374 |
|  |  | XRCC4 | 7518 |

**Supplemental Table 5.**
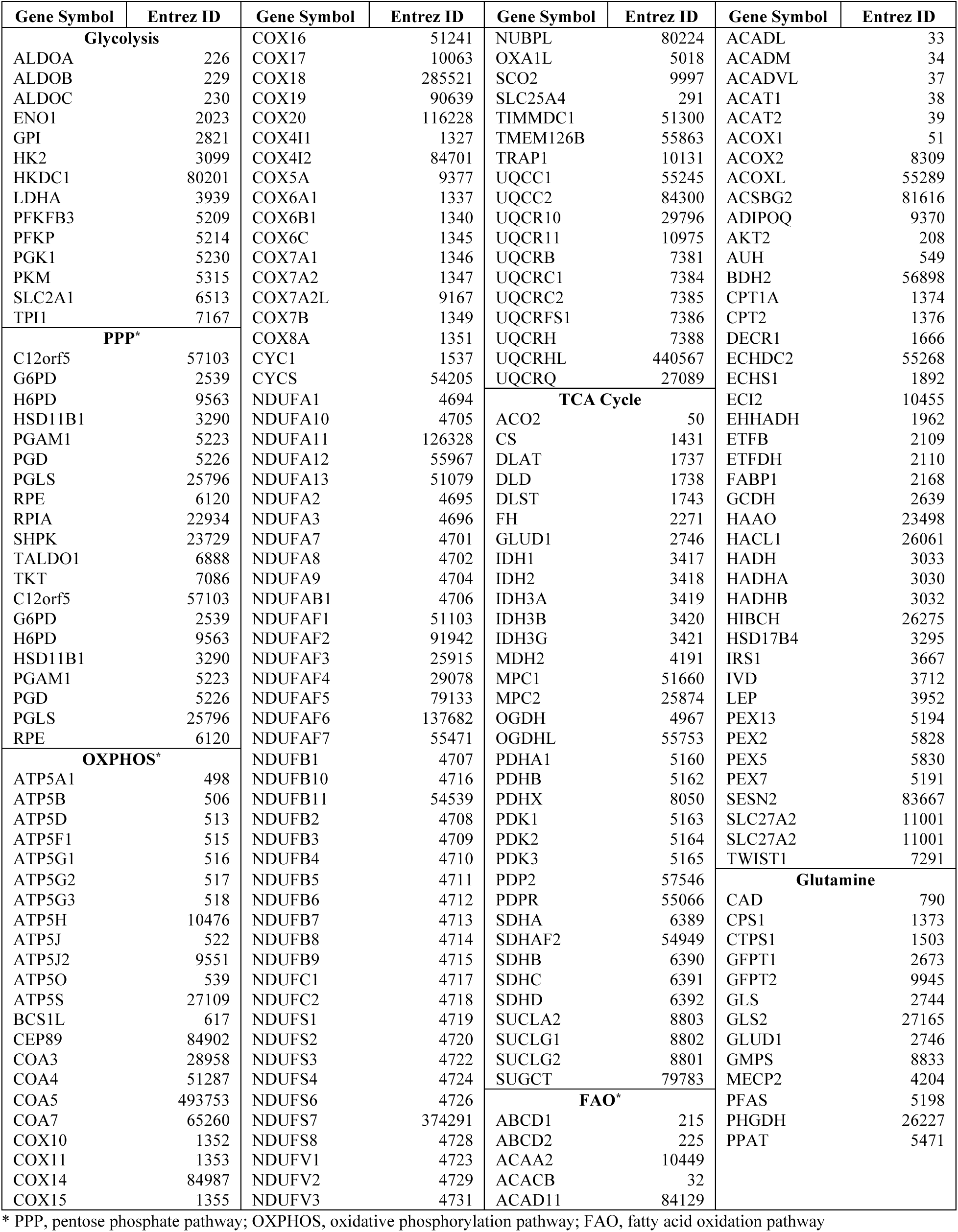
Input list of human gene sets of metabolic pathways for Gene Set Variation Analysis.

## REFERENCES

1. Nutt SL, Tarlinton DM. Germinal center B and follicular helper T cells: siblings, cousins or just good friends? Nat Immunol. 2011;12(6):472–477.

2. Domeier PP, Schell SL, Rahman ZS. Spontaneous germinal centers and autoimmunity. Autoimmunity. 2017;50(1):4–18.

3. Crow MK. Type I interferon in the pathogenesis of lupus. J Immunol. 2014;192(12):5459–5468.

4. Tsokos GC. Autoimmunity and organ damage in systemic lupus erythematosus. Nat Immunol. 2020;21(6):605–614.

5. Kim J, Gross JA, Dillon SR, Min JK, Elkon KB. Increased BCMA expression in lupus marks activated B cells, and BCMA receptor engagement enhances the response to TLR9 stimulation. Autoimmunity. 2011;44(2):69–81.

6. Salazar-Camarena DC, Palafox-Sanchez CA, Cruz A, Marin-Rosales M, Munoz-Valle JF. Analysis of the receptor BCMA as a biomarker in systemic lupus erythematosus patients. Sci Rep. 2020;10(1):6236.

7. Navarra SV, Guzman RM, Gallacher AE, et al. Efficacy and safety of belimumab in patients with active systemic lupus erythematosus: a randomised, placebo-controlled, phase 3 trial. Lancet. 2011;377(9767):721–731.

8. Zhang X, Lindwall E, Gauthier C, et al. Circulating CXCR5+CD4+helper T cells in systemic lupus erythematosus patients share phenotypic properties with germinal center follicular helper T cells and promote antibody production. Lupus. 2015;24(9):909–917.

9. Ise W, Fujii K, Shiroguchi K, et al. T Follicular Helper Cell-Germinal Center B Cell Interaction Strength Regulates Entry into Plasma Cell or Recycling Germinal Center Cell Fate. Immunity. 2018;48(4):702–715 e704.

10. Dolff S, Abdulahad WH, Westra J, et al. Increase in IL-21 producing T-cells in patients with systemic lupus erythematosus. Arthritis Res Ther. 2011;13(5):R157.

11. Terrier B, Costedoat-Chalumeau N, Garrido M, et al. Interleukin 21 correlates with T cell and B cell subset alterations in systemic lupus erythematosus. J Rheumatol. 2012;39(9):1819–1828.

12. Gigoux M, Shang J, Pak Y, et al. Inducible costimulator promotes helper T-cell differentiation through phosphoinositide 3-kinase. Proc Natl Acad Sci U S A. 2009;106(48):20371–20376.

13. Pearce EL, Pearce EJ. Metabolic pathways in immune cell activation and quiescence. Immunity. 2013;38(4):633–643.

14. Chang CH, Curtis JD, Maggi LB, Jr., et al. Posttranscriptional control of T cell effector function by aerobic glycolysis. Cell. 2013;153(6):1239–1251.

15. Caro-Maldonado A, Wang R, Nichols AG, et al. Metabolic reprogramming is required for antibody production that is suppressed in anergic but exaggerated in chronically BAFF-exposed B cells. J Immunol. 2014;192(8):3626–3636.

16. Yin Y, Choi SC, Xu Z, et al. Normalization of CD4+ T cell metabolism reverses lupus. Sci Transl Med. 2015;7(274):274ra218.

17. Choi SC, Titov AA, Abboud G, et al. Inhibition of glucose metabolism selectively targets autoreactive follicular helper T cells. Nat Commun. 2018;9(1):4369.

18. McPhee CG, Sproule TJ, Shin DM, et al. MHC class I family proteins retard systemic lupus erythematosus autoimmunity and B cell lymphomagenesis. J Immunol. 2011;187(9):4695–4704.

19. Laszlo G, Hathcock KS, Dickler HB, Hodes RJ. Characterization of a novel cell-surface molecule expressed on subpopulations of activated T and B cells. J Immunol. 1993;150(12):5252–5262.

20. Cervenak L, Magyar A, Boja R, Laszlo G. Differential expression of GL7 activation antigen on bone marrow B cell subpopulations and peripheral B cells. Immunol Lett. 2001;78(2):89–96.

21. Baumjohann D, Preite S, Reboldi A, et al. Persistent antigen and germinal center B cells sustain T follicular helper cell responses and phenotype. Immunity. 2013;38(3):596–605.

22. Suarez-Fueyo A, Bradley SJ, Tsokos GC. T cells in Systemic Lupus Erythematosus. Curr Opin Immunol. 2016;43:32–38.

23. Catalina MD, Bachali P, Yeo AE, et al. Patient ancestry significantly contributes to molecular heterogeneity of systemic lupus erythematosus. JCI Insight. 2020;5(15).

24. Mootha VK, Lindgren CM, Eriksson KF, et al. PGC-1alpha-responsive genes involved in oxidative phosphorylation are coordinately downregulated in human diabetes. Nat Genet. 2003;34(3):267–273.

25. Subramanian A, Tamayo P, Mootha VK, et al. Gene set enrichment analysis: a knowledge-based approach for interpreting genome-wide expression profiles. Proc Natl Acad Sci U S A. 2005;102(43):15545–15550.

26. Hanzelmann S, Castelo R, Guinney J. GSVA: gene set variation analysis for microarray and RNA-seq data. BMC Bioinformatics. 2013;14:7.

27. Bengsch B, Johnson AL, Kurachi M, et al. Bioenergetic Insufficiencies Due to Metabolic Alterations Regulated by the Inhibitory Receptor PD-1 Are an Early Driver of CD8(+) T Cell Exhaustion. Immunity. 2016;45(2):358–373.

28. Qie S, Yoshida A, Parnham S, et al. Targeting glutamine-addiction and overcoming CDK4/6 inhibitor resistance in human esophageal squamous cell carcinoma. Nat Commun. 2019;10(1):1296.

29. Weisel FJ, Mullett SJ, Elsner RA, et al. Germinal center B cells selectively oxidize fatty acids for energy while conducting minimal glycolysis. Nat Immunol. 2020;21(3):331–342.

30. Li W, Titov AA, Morel L. An update on lupus animal models. Curr Opin Rheumatol. 2017;29(5):434–441.

31. Sitrin J, Suto E, Wuster A, et al. The Ox40/Ox40 Ligand Pathway Promotes Pathogenic Th Cell Responses, Plasmablast Accumulation, and Lupus Nephritis in NZB/W F1 Mice. J Immunol. 2017;199(4):1238–1249.

32. Aft RL, Zhang FW, Gius D. Evaluation of 2-deoxy-D-glucose as a chemotherapeutic agent: mechanism of cell death. Br J Cancer. 2002;87(7):805–812.

33. Song M, Sandoval TA, Chae CS, et al. IRE1alpha-XBP1 controls T cell function in ovarian cancer by regulating mitochondrial activity. Nature. 2018;562(7727):423–428.

34. van Schadewijk A, van’t Wout EF, Stolk J, Hiemstra PS. A quantitative method for detection of spliced X-box binding protein-1 (XBP1) mRNA as a measure of endoplasmic reticulum (ER) stress. Cell Stress Chaperones. 2012;17(2):275–279.

35. Spitz DR, Sim JE, Ridnour LA, Galoforo SS, Lee YJ. Glucose deprivation-induced oxidative stress in human tumor cells. A fundamental defect in metabolism? Ann N Y Acad Sci. 2000;899:349–362.

36. Simons AL, Mattson DM, Dornfeld K, Spitz DR. Glucose deprivation-induced metabolic oxidative stress and cancer therapy. J Cancer Res Ther. 2009;5 Suppl 1:S2–6.

37. Dent AL, Shaffer AL, Yu X, Allman D, Staudt LM. Control of inflammation, cytokine expression, and germinal center formation by BCL-6. Science. 1997;276(5312):589–592.

38. Nurieva RI, Chung Y, Martinez GJ, et al. Bcl6 mediates the development of T follicular helper cells. Science. 2009;325(5943):1001–1005.

39. Schmidts A, Ormhoj M, Choi BD, et al. Rational design of a trimeric APRIL-based CAR-binding domain enables efficient targeting of multiple myeloma. Blood Adv. 2019;3(21):3248–3260.

40. Rhoads JP, Major AS, Rathmell JC. Fine tuning of immunometabolism for the treatment of rheumatic diseases. Nat Rev Rheumatol. 2017;13(5):313–320.

41. Perl A. Activation of mTOR (mechanistic target of rapamycin) in rheumatic diseases. Nat Rev Rheumatol. 2016;12(3):169–182.

42. Teng X, Cornaby C, Li W, Morel L. Metabolic regulation of pathogenic autoimmunity: therapeutic targeting. Curr Opin Immunol. 2019;61:10–16.

43. Galgani M, Bruzzaniti S, Matarese G. Immunometabolism and autoimmunity. Curr Opin Immunol. 2020;67:10–17.

44. Mazumdar C, Driggers EM, Turka LA. The Untapped Opportunity and Challenge of Immunometabolism: A New Paradigm for Drug Discovery. Cell Metab. 2020;31(1):26–34.

45. Brand KA, Hermfisse U. Aerobic glycolysis by proliferating cells: a protective strategy against reactive oxygen species. FASEB J. 1997;11(5):388–395.

46. Jiang C, Loo WM, Greenley EJ, Tung KS, Erickson LD. B cell maturation antigen deficiency exacerbates lymphoproliferation and autoimmunity in murine lupus. J Immunol. 2011;186(11):6136–6147.

47. O’Connor BP, Raman VS, Erickson LD, et al. BCMA is essential for the survival of long-lived bone marrow plasma cells. J Exp Med. 2004;199(1):91–98.

48. Peperzak V, Vikstrom I, Walker J, et al. Mcl-1 is essential for the survival of plasma cells. Nat Immunol. 2013;14(3):290–297.

49. Pradelli LA, Beneteau M, Chauvin C, et al. Glycolysis inhibition sensitizes tumor cells to death receptors-induced apoptosis by AMP kinase activation leading to Mcl-1 block in translation. Oncogene. 2010;29(11):1641–1652.

50. Raez LE, Papadopoulos K, Ricart AD, et al. A phase I dose-escalation trial of 2-deoxy-D-glucose alone or combined with docetaxel in patients with advanced solid tumors. Cancer Chemother Pharmacol. 2013;71(2):523–530.

51. Bhatt AN, Shenoy S, Munjal S, et al. 2-Deoxy-D-Glucose as an Adjunct to Standard of Care in the Medical Management of COVID-19: A Proof-of-Concept & Dose-Ranging Randomised Clinical Trial. medRxiv. 2021:doi: https://doi.org/10.1101/2021.1110.1108.21258621.

52. Stein M, Lin H, Jeyamohan C, et al. Targeting tumor metabolism with 2-deoxyglucose in patients with castrate-resistant prostate cancer and advanced malignancies. Prostate. 2010;70(13):1388–1394.

53. Hemmi H, Kaisho T, Takeuchi O, et al. Small anti-viral compounds activate immune cells via the TLR7 MyD88-dependent signaling pathway. Nat Immunol. 2002;3(2):196–200.

54. Li B, Dewey CN. RSEM: accurate transcript quantification from RNA-Seq data with or without a reference genome. BMC Bioinformatics. 2011;12:323.

55. Langmead B, Trapnell C, Pop M, Salzberg SL. Ultrafast and memory-efficient alignment of short DNA sequences to the human genome. Genome Biol. 2009;10(3):R25.

56. Robinson MD, McCarthy DJ, Smyth GK. edgeR: a Bioconductor package for differential expression analysis of digital gene expression data. Bioinformatics. 2010;26(1):139–140.

57. Bult CJ, Blake JA, Smith CL, Kadin JA, Richardson JE, Mouse Genome Database G. Mouse Genome Database (MGD) 2019. Nucleic Acids Res. 2019;47(D1):D801–D806.

58. Heng TS, Painter MW, Immunological Genome Project C. The Immunological Genome Project: networks of gene expression in immune cells. Nat Immunol. 2008;9(10):1091–1094.

59. Kingsmore KM, Bachali P, Catalina MD, et al. Altered expression of genes controlling metabolism characterizes the tissue response to immune injury in lupus. Sci Rep. 2021;11(1):14789.

60. Andreoletti G, Lanata CM, Trupin L, et al. Transcriptomic analysis of immune cells in a multi-ethnic cohort of systemic lupus erythematosus patients identifies ethnicity- and disease-specific expression signatures. Commun Biol. 2021;4(1):488.

